# Temporal variability and cell mechanics control robustness in mammalian embryogenesis

**DOI:** 10.1101/2023.01.24.525420

**Authors:** Dimitri Fabrèges, Bernat Corominas Murtra, Prachiti Moghe, Alison Kickuth, Takafumi Ichikawa, Chizuru Iwatani, Tomoyuki Tsukiyama, Nathalie Daniel, Julie Gering, Anniek Stokkermans, Adrian Wolny, Anna Kreshuk, Véronique Duranthon, Virginie Uhlmann, Edouard Hannezo, Takashi Hiiragi

## Abstract

How living systems achieve precision in form and function despite their intrinsic stochasticity is a fundamental yet open question in biology. Here, we establish a quantitative morphomap of pre-implantation embryogenesis in mouse, rabbit and monkey embryos, which reveals that although blastomere divisions desynchronise passively without compensation, 8-cell embryos still display robust 3D structure. Using topological analysis and genetic perturbations in mouse, we show that embryos progressively change their cellular connectivity to a preferred topology, which can be predicted by a simple physical model where noise and actomyosin-driven compaction facilitate topological transitions lowering surface energy. This favours the most compact embryo packing at the 8- and 16-cell stage, thus promoting higher number of inner cells. Impairing mitotic desynchronisation reduces embryo packing compactness and generates significantly more cell mis-allocation and a lower proportion of inner-cell-mass-fated cells, suggesting that stochasticity in division timing contributes to achieving robust patterning and morphogenesis.

## Introduction

Living systems rely on molecular and cellular mechanisms with intrinsically stochastic dynamics. Nonetheless, they establish robust forms and functions on multiple scales with complex interplay between molecules, cells, tissues and species across evolutionary time. Cell organisation and decision making have been largely considered instructive, i.e., a signal instructs the recipient cell to differentiate or trigger some signalling pathway (Chubb, 2017), which may lead to the notion that variability is destructive and must be reduced or filtered out. However, it remains unknown whether robust mechanisms and decision making have been evolutionarily selected to accommodate stochasticity, or if the processes at the source of variability have been selected to grant robustness. The last decades have shown a growing interest for a decision making paradigm based on stochastic dynamics (Balázsi et al., 2011; Eldar and Elowitz, 2010; Raj and van Oudenaarden, 2008; Spratt and Lane, 2022), particularly in bacteria where gene expression variability was successfully manipulated to demonstrate its role in cell differentiation (Maamar et al., 2007; Süel et al., 2007). Stochastic processes have since been shown to be involved in a wide variety of context and species, including cell fate specification (Chang et al., 2008; Lord et al., 2019; Maamar et al., 2007; Meyer and Roeder, 2014; Ohnishi et al., 2014; Simon et al., 2018; Süel et al., 2007), cancer adaptation (Bell and Gilan, 2020; Brock et al., 2009; Feinberg and Irizarry, 2010; Feinberg et al., 2006; Koldobskiy et al., 2021), embryo morphogenesis (Carlson et al., 2015; Dumollard et al., 2017), leaf formation (Hong et al., 2016; Hong et al., 2018), tissue folding (Haas et al., 2018), cell sorting (Yanagida et al., 2022) and evolvability (Bromham, 2003; Draghi, 2019; Hernández et al., 2022; Schmid et al., 2022).

Early mammalian embryos have been shown to exhibit intra- and inter-embryo variability in gene expression (Ohnishi et al., 2014; Dietrich and Hiiragi, 2007; Roberts et al., 2011; Lavagi et al., 2018), making them an excellent model to study the role of variability in a minimal and robust system. Not only gene expression, but other mechanical and temporal parameters exhibit measurable variability within and between embryos that may affect patterning and morphogenesis. For instance, the 2^nd^ cleavage orientation was shown to be random (Louvet-Vallée et al., 2005); the variability in cleavage timing has been suggested to impact cell-fate segregation (Kelly et al., 1978; Mashiko et al., 2022); the nuclear-cytoplasmic ratio has been linked to cell differentiation (Aiken et al., 2004); and heterogeneity in cell contractility was shown to drive cell sorting in 16-cell stage mouse embryo (Maître et al., 2016; Samarage et al., 2015) while cell-to-cell variability in cellular fluidity may contribute to the segregation of primitive endoderm and epiblast in blastocysts (Yanagida et al., 2022). Although recent studies in mouse showed that at least some of these variabilities are regulated by the feedbacks between cell polarity, tissue mechanics and gene expression (Maître et al., 2016; Korotkevich et al., 2017), the role that intercellular variabilities may play in development and its robustness has not been explored yet.

To formally address the role of variability, it needs to be quantified with an adequate number of samples suitable for statistical analyses, and tested with its manipulation in space and/or time. In this study, we thus developed such an experimental system using mouse pre-implantation embryos. While spatial organisation of development is relatively well studied in the context of morphogenesis and patterning, less is known about the temporal regulation of developmental progression. Therefore, we started characterising and measuring the variability among cells in developmental timing.

## Results

### Embryo variability in cleavage timing increases at a constant and species-specific rate

To characterise the variability of the developmental timing of pre-implantation mouse embryos, we first measured the natural variability in cleavage timing between cells within an embryo (Figure 1A and Video S1). The cell cycles and cleavages of mammalian embryos run asynchronously among blastomeres (Bowman and McLaren) but their variability and potential correlation have not been quantitatively characterised. The duration of the 3^rd^ cleavage (4- to 8-cell stage), for example, ranged from 30 minutes to 3h30 depending on the embryo, accompanied by variable duration of the intermitotic period (Pearson correlation *R* = −0.606 (*P* < 0.05), Figure S1A). To assess whether the timing of divisions is actively regulated or coordinated among cells, we built a semi-automatic cell-tracking pipeline and quantified cleavage timing in 16 embryos, from the 4-to the 32-or 64-cell stage (Figure 1B,C). For this, we tested the null-hypothesis that cells do not show any coordination: assuming the cell cycle length follows a normally distributed random variable of variance 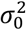, the successive accumulation of mitosis timing differences should be normally distributed, with variance linearly increasing with the cleavage number (Figure 1D). Deviation from the linear relationship between variance and cleavage number would be indicative of an active synchronisation or desynchronisation by the cell’s nearby environment (Figure S1B). In line with the null hypothesis, the distribution of the division timing measured for the 3^rd^, 4^th^, 5^th^ and 6^th^ cleavage followed a normal distribution (Figure 1E) and linearly correlated with the cleavage number (Pearson correlation *R* = 0.522 (*P* < 10^−7^), Figure 1F). Although the average cell cycle length increased after the 5^th^ cleavage (+1 hour, Figure S1C), due to a longer cell cycle for cells in the inner cell mass (ICM) compared to cells in the trophectoderm (TE, 1.5 hour of difference, Figure S1D), we did not find significant correlation between cell position and timing of division, suggesting that the lengthening of the cell cycle was due to cell differentiation and not cell-cell interaction (Figure 1G). To confirm this, we experimentally introduced a substantial asynchrony in the cleavage timing by inserting a time-shifted 1/8^th^ blastomere under the zona pellucida of host embryos at the 8-cell stage, effectively generating heterochronic chimera (Figure 1H). However, neither the time-shift of the donor cell or that of the host cell was affected during the following cleavages (Figure 1I). Altogether, these observations indicate a lack of discernible cell-cell coordination in cleavage timing, which results in a cell-autonomous desynchronisation at a constant rate.

**Figure 1.**
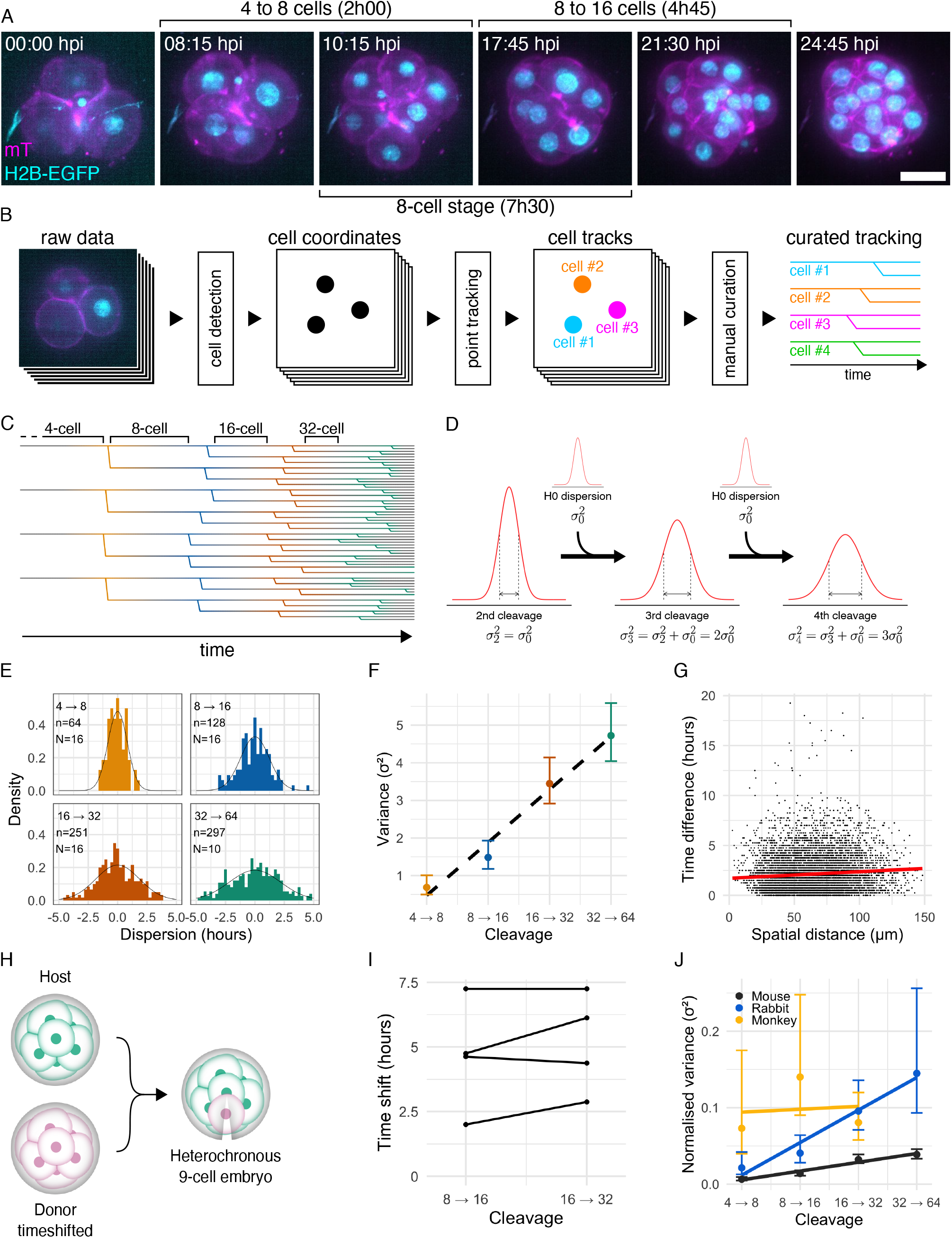
Embryo variability in cleavage timing increases at a constant and species-specific rate. (**A**) Maximum intensity projection of a representative live imaging dataset, out of 16 datasets from 5 independent experiments, of a mouse embryo expressing mT (magenta); H2B-EGFP (cyan) developing from the 4-to the 16-cell stage. Time post imaging (hh:mm); Scale bar, 25 *µ*m. See also Video S1. (**B**) Schematic representation of the tracking pipeline. 3D+time microscopy data were analysed to automatically detect nuclei centre position, then automatically tracked and manually curated to ensure maximum accuracy. See also Methods. (**C**) Representative curated tracking displaying the 3rd (yellow), 4th (blue), 5th (red) and 6th (green) cleavages covering a period of 43 hours from left to right. Each branch represents a cell. Each branching represents a mitosis. (**D**) Schematic representation of the null hypothesis (H0) under which the distribution of the timing of mitosis at the n^th^ cleavage is normally distributed with a variance of 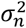 and equals the cumulative sum of independent normally distributed random variables with a variance of 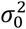 such that 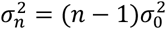. See also Figure S2B. (**E**) Density distribution of the timing of mitosis around the mean grouped by cleavage number, measured from 16 embryos of 5 independent experiments. Black line, gaussian fit of each distribution (*s. d*. = 0.83, 1.22, 1.86 and 2.17 for the 3^rd^, 4^th^, 5^th^ and 6^th^ cleavage respectively). n, number of mitoses. N, number of embryos. Colour code as in (C). (**F**) Variance in mitosis timing as a function of cleavage number. Dashed line, linear regression. Pearson correlation *R* = 0.989 (*P* = 0.011, *CI*_95%_ = [0.552; 0.999]). Colour code as in (C). (**G**) Time difference in mitosis timing as a function of spatial distance for each pair of same-generation cells (black dots), measured for 7856 pairs of cells in 16 embryos from 5 independent experiments. Red line, linear regression. Pearson correlation *R* = 0.085 (*P* < 10^−13^, *CI*_95%_ = [0.063; 0.107]). (**H**) Diagram of the experimental generation of heterochronic chimera. A time-shifted blastomere from an 8-cell stage donor (magenta) is inserted under the zona pellucida of 8-cell stage host (cyan), generating a 9 cells heterochronic chimera. See also Methods. (**I**) Evolution of the shift in timing of division between the 4^th^ and the 5^th^ cleavage between host and donor in 9-cell heterochronic embryos. (**J**) Comparison of the variance of the mitosis timing normalised by the cell cycle length, as a function of the cleavage number in mouse (black, 16 embryos from 5 independent experiments), rabbit (blue, 6 embryos from 3 independent experiments) and monkey (yellow, 4 embryos from 2 independent experiments). Thick lines, linear regression. Error bars, 95% confidence interval.

Since various mammalian embryos undergo asynchronous cleavage cycles, we performed a similar characterisation of variability in division timing by injecting rabbit and monkey embryos with mRNAs encoding myrTagRFP-T;H2B-EGFP and mT-T2A-H2B-EGFP, respectively (Figure S1E,F and Video S2 and S3). Live-imaging microscopy and its analysis showed that rabbit and monkey embryos share a similar desynchronisation pattern, with their desynchronisation rate higher than that of mouse embryos (Figure 1J). Notably, the desynchronisation rate is species-specific, which suggests that variability in cleavage timing may be an evolutionary trait that plays a role in the following developmental processes.

### Embryo spatial variability reduces during the 8-cell stage

To investigate the impact of cleavage timing variability on the robustness of development, we developed a pipeline to parameterise cell shape and build a statistical vector map of morphogenesis – or morphomap (Figure 2A). First, cells were automatically labelled, followed by manual curation, from 3D-images of H2B-GFP; mT transgenic embryos. Then, the surface of the labelled objects was fitted with exponential splines (Delgado-Gonzalo et al., 2013) (see Figure S2A,B and Methods for details), generating a set of 63 parameters describing both the shape of each cell and its relative position in the embryo. Each embryo was thus geometrically described by a unique set of 63 parameters times 8 blastomeres: 504 parameters, projected in 2D for visualisation. The tSNE 2D-projection of the 504D-morphomap of 29 embryos (Figure 2B) and measurement of pair-wise 446,985 distances between 946 3D-images of embryos developing during the 8-cell stage revealed that embryos become more and more geometrically similar and converge towards a specific area of the morphomap (Pearson correlation *R* = −0.228 (*P* < 10^−15^), Figures 2C and S2C).

**Figure 2.**
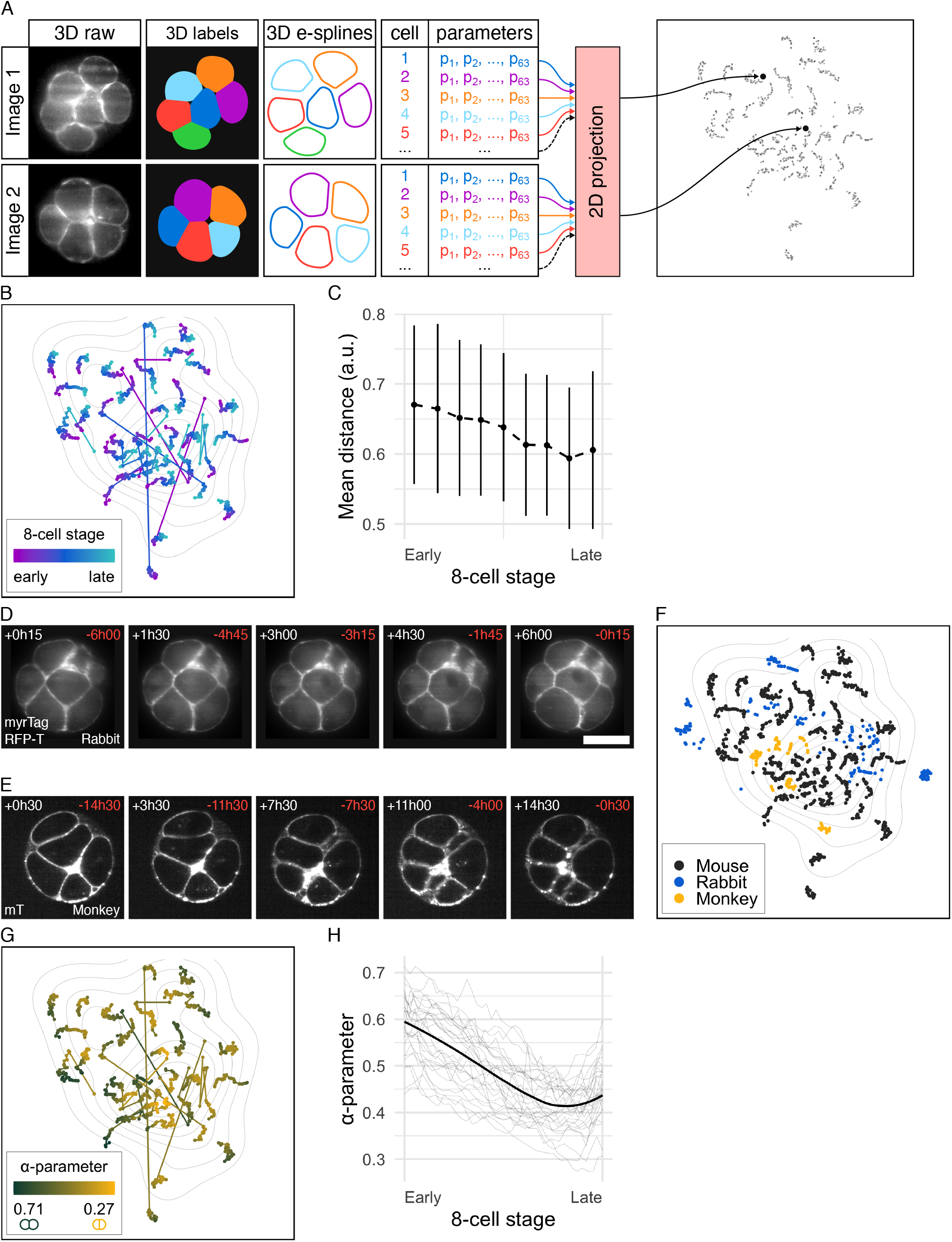
Embryo spatial variability reduces during the 8-cell stage. (**A**) Descriptive representation of the geometrical distance measurement pipeline. 3D images of embryo membranes (first column) were automatically segmented, manually curated to generate high accuracy labelled volumes (second column), and fitted with exponential splines (third column) producing a set of 504 parameters for each embryo at each timepoint (fourth column). The distance between each pair of 3D images was computed from the exponential spline parameters, and used to project the 504D-morphomap on a 2D plane (see also Methods). (**B,F,G**) tSNE projection of the morphomap. Each embryo is represented with a sequence timepoints and coloured as a function of the normalised progression though the 8-cell stage (B, early in magenta to late in cyan), the species (F, mouse in black, rabbit in blue and monkey in yellow) or the compaction parameter (G, high alpha or low compaction in green, low alpha or high compaction in yellow). Isolines, density map of the end of the 8-cell stage. *n* = 29 mouse embryos, 10 rabbit embryos and 4 monkey embryos. (**C,H**) Normalised time course through the 8-cell stage of the mean ± s.d. of the 406 pair-wise geometrical distances (C) and the mean compaction parameter *α* (H) in 29 embryos. Light grey lines, individual embryo tracks. See also Figure S2E,F. (**D,E**) Cross-section of a representative live imaging dataset of a rabbit embryo (D, *n* = 10 embryos from 4 independent experiments) and a monkey embryo (E, *n* = 4 embryos from 2 independent experiments) expressing myrTagRFP-T (D) and mT (E). Scale bar, 50 *µ*m. Time after the beginning (white, top-left corner) and before the end (red, top-right corner) of the 8-cell stage. See also Figure S2B,C.

Likewise, we built the morphomap for rabbit (*n* = 10, Figures 2D and S2D) and monkey (*n* = 4, Figures 2E and S2E) embryos to examine whether their embryos show similar geometrical convergence (Figure 2F). Rabbit and monkey embryos were larger in volume than mouse embryos (Figure S2F), hence their total volume was normalised to allow for direct comparison. Remarkably, the three species exhibited similar geometrical structures and shared the same morphomap. The absence of cluster by species suggests that similar design principles could govern embryo shape changes during the 8-cell stage in the three species.

In search of such a principle driving geometrical convergence, we looked into compaction. In mouse embryos, compaction starts at the 8-cell stage which results in significant cell shape changes and an overall smoothing of embryo surface. Using the compaction parameter *α* (defined as the ratio between cell-cell and cell-bulk surface tensions (Maître et al., 2015), Figure S2G), we quantified the compaction over time and showed a significant decrease of the *α*-parameter, indicative of an increase of the degree of compaction and cell shape change (Figures 2G,H and S2G). However, these changes by compaction were not sufficient to explain the geometrical convergence, since the decrease in inter-embryo distance correlates with *α* when it is below 0.4 (Figure S2H), while many embryos did not reach *α* below 0.4 (Figure 2H). This suggests that cell shape changes due to compaction are not sufficient to explain the observed geometrical convergence.

### Embryo topological variability reduces during the 8-cell stage

We reasoned that the morphomap encompasses both shape and arrangement of the cells. While the former is linked to the geometrical changes induced by compaction, the latter depends on the cell-cell contact structure, that is, in the topological properties of the embryo packing (Giammona and Campàs, 2021; Imran Alsous et al., 2018; Kuang et al., 2022; Petridou et al., 2021). Therefore, to identify the mechanism driving morphological convergence, we examined cellular topology in the 8-cell-stage embryo. For 8 adhesive passive spheres, it has been shown that although there is an infinitely large number of 3D geometrical arrangements, only 13 packings are rigid, i.e., correspond to a local energy minimum where the configuration is stable as any relative cell-cell displacement costs finite energy (Arkus et al., 2009; Arkus et al., 2011; Jacobs, 1998) (Figures 3A and S3A). We thus used these 13 packings as landmarks to interpret our morphomaps, and classified embryos either as non-rigid (NR) or as belonging to one of the rigid packings, based on the topological proximity of the cell-cell contact structure of the embryo to one of the 13 rigid packings (see Methods for details). This analysis revealed that although many embryos were non-rigid at the start of the 8-cell stage, nearly all of them converged towards one of the 13 rigid packings at the end (Figure S3B). Strikingly, we could observe an additional and unexpected topological convergence within rigid packings, as two specific rigid packings, D_2d_ and C_s_(2), became highly overly represented over time (30.3% and 29.8%, respectively, of 29 embryos, Figure S3B). All the other rigid packings were grouped and referred to as “Others” in subsequent analyses. Remarkably, D_2d_ overlaps with the attractor region detected by our morphospace analysis (Figure 3B). However, the inter-embryo distance within all embryo packings was stable over the course of the 8-cell stage (Figure 3C), suggesting that the overall geometrical convergence is linked to topological transitions from geometrically heterogeneous and NR packings to the most homogeneous one (D_2d_). To test this prediction, we analysed the evolution of the topological transitions over time. Fifty-three topological transitions were observed from 29 embryos during which cell contacts were created, lost or strengthened (Figure 3D, Video S4). Indeed, transitions from NR to Others, Others to Others, Others to C_s_(2) and C_s_(2) to D_2d_ were overly represented (Figure 3E) resulting in the significant decrease of non-rigid packings described earlier and in a dramatic increase of D_2d_ proportion over the course of the 8-cell stage (Figure 3F). A similar topological transition was also observed in rabbit and monkey embryos (Figure 3G). Collectively, these analyses show that the overall geometrical convergence among mammalian embryos is driven by a chain of topological transitions towards D_2d_ via C_s_(2).

**Figure 3.**
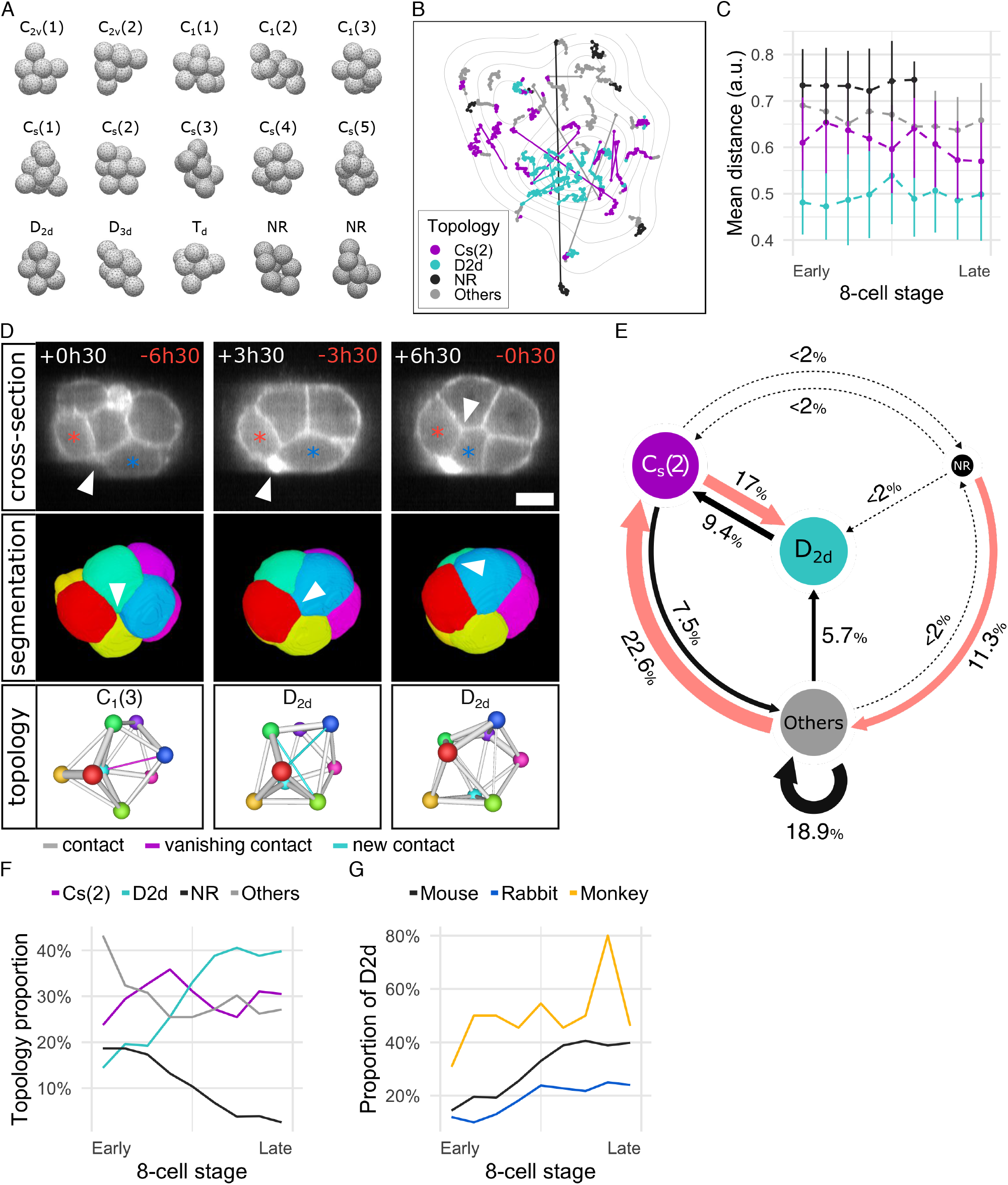
Embryo topological variability reduces during the 8-cell stage. (**A**) The 13 rigid packings and two examples of non-rigid packings (NR) for clusters of 8 spheres. Rigid packings were named using the Schoenflies notation and describes the symmetry of the packing in three dimensions. A number between parenthesis was added to differentiate packings sharing the same Schoenflies notation. See also Figure S3A. (**B**) tSNE projection of the morphomap. Each embryo is represented with a sequence of connected timepoints and coloured as a function of the topological proximity to C_s_(2) (magenta), D_2d_ (cyan), or any other rigid packing (grey). Non rigid packings (NR) are coloured in black. Isolines, density map of the end of the 8-cell stage. *n* = 29 mouse embryos. (**C**) Normalised time course through the 8-cell stage of the mean ± s.d. of pair-wise geometrical distances within embryos categorized and coloured as in (B). (**D**) Cross-section (first row), surface of cell segmentation (second row) and topological network (third row) of a representative topological transition from C_1_(3) (first column) to D_2d_ (second and third columns). Arrow heads point the absence of cell-cell contact (first column), the initiation of a contact (second column), and the expansion of the contact (third column) between two cells respectively marked with a red and blue star (first row) or represented in red and blue (second and third row). Cell-cell contact loss (magenta) and gain (cyan) has been highlighted on the topology network among lasting contacts (grey). Time after the beginning (white, top-left corner) and before the end (red, top-right corner) of the 8-cell stage. Scale bar, 25 *µ*m. See also Video S4. (**E**) Topological transition map as observed in mouse embryos at the 8-cell stage between C_s_(2) (magenta), D_2d_ (cyan), non-rigid (NR, black) or any other rigid packings (Others, grey). An arrow indicates that a topological transition between two groups was observed once (dotted lines) or more (solid lines). Circle area is proportional to the topology proportion. Arrows are labelled and sized by the occurrence of the transition. Red arrows indicate the direction of the net flux of transitions. *n* = 53 transitions. (**F**) Evolution of the proportion of C_s_(2) (magenta), D_2d_ (cyan), non-rigid (NR, black) or any other rigid packings (Others, grey) as a function of the normalised progression through the 8-cell stage. *n* = 29 mouse embryos. See also Figure S2B. (**G**) Evolution of the proportion of topologies identified as D_2d_ in mouse (black, *n* = 29 embryos), rabbit (blue, *n* = 10 embryos) and monkey (yellow, *n* = 4 embryos) as a function of the normalised progression through the 8-cell stage.

### Surface energy minimisation with compaction is sufficient to recapitulate geometrical and topological convergence

This convergence towards a single packing was surprising as it was shown for attractive hard spheres that all 13 rigid packings have the same energy (Arkus et al., 2009). Upon compaction, cell-cell adhesion configuration has been shown to be well-described by a soap-bubble model (Goel et al., 1986; Hayashi and Carthew, 2004; Heisenberg, 2017; Maître et al., 2015; Maître et al., 2016; Pierre et al., 2016) where the relative energy can be determined with the cell-cell contact surface area, the contact-free surface area (not in contact with other cells) and the relative surface tension or *α*-parameter (Figure 4A, see also Methods). To test how cell-cell adhesion and compaction (given *α*-parameter) changes the stabilities of embryo packings, we thus used the Surface Evolver software (Brakke, 1992) to simulate morphogenesis – in both a theoretical and data-driven manner. Using the 13 rigid packings (Figure 3A) as initial conditions to our model, we simulated the compaction from low to high compaction (*α* = 0.8 and 0.3 respectively, Figures 4B,C and S4A,B). Crucially, compaction leads to few specific packings having a lower energy than others (with D_2d_ being the lowest, Figure 4D), and as a consequence, the model predicted a dramatic convergence at high compaction towards D_2d_ for most of the rigid packings (Figures 4E,F and S4C). Several of these transitions could occur even with low level of noise, due to compaction near-deterministically creating new cell contacts (Figure S4D,E) while a number of others required cell re-arrangements (or T1 transitions), which has been shown to be associated to energy barriers across different modelling frameworks (Bi et al., 2014; Bi et al., 2016; Marmottant et al., 2009; Pinheiro et al., 2022), and thus noise to go from one minimum to another (Figure 4C, see also Methods). We also found that C_s_(2) was the second most stable packing, functioning as a transition intermediary towards D_2d_ in the simulations. The model thus closely mirrored our experimental datasets for topological transitions observed in 8-cell stage embryo.

**Figure 4.**
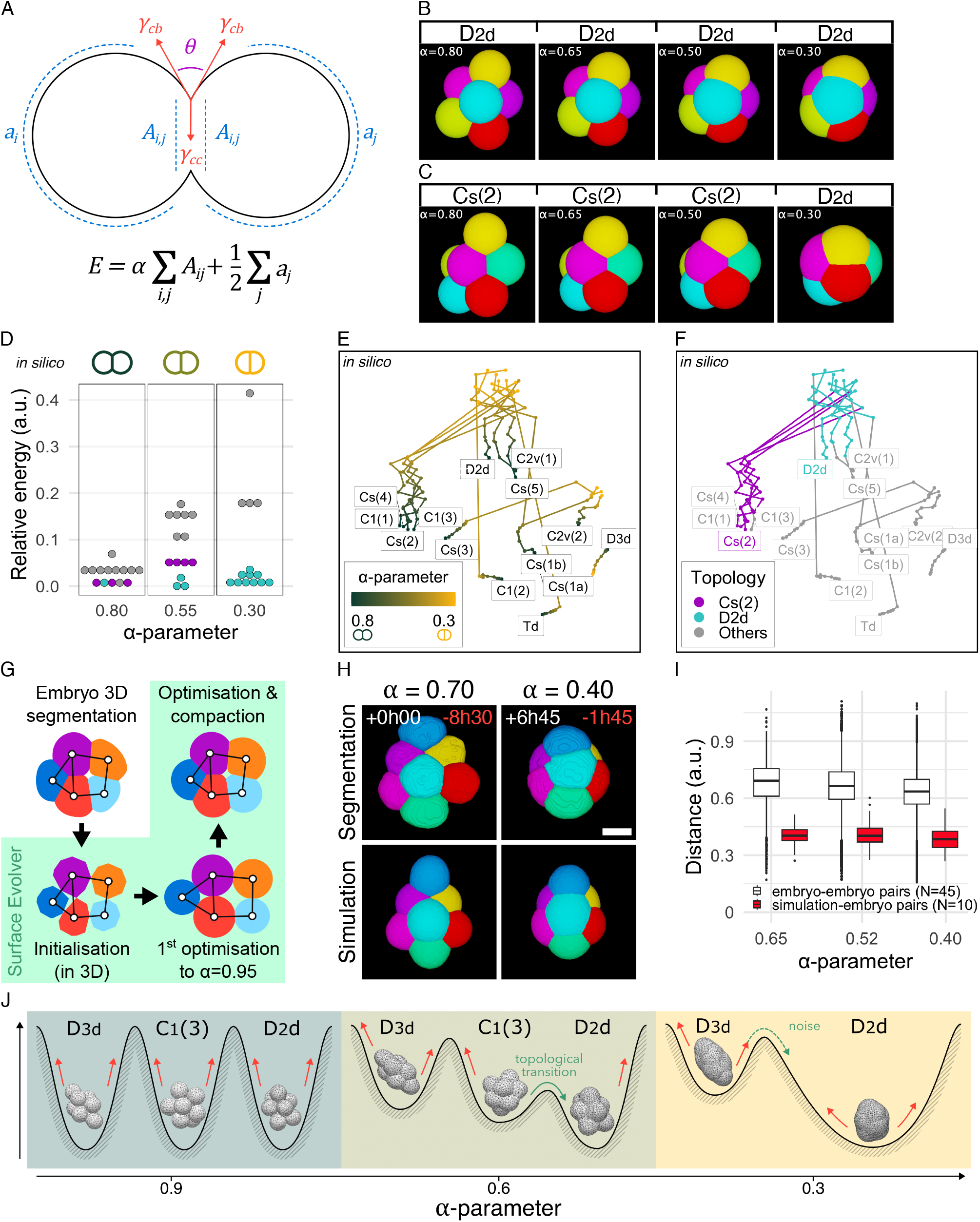
Surface energy minimisation with compaction is sufficient to recapitulate geometrical and topological convergence. (**A**) Diagram of the biophysical model. The total energy of an embryo depends on cell-cell contact area (*A*_*i,j*_ between cells *i* and *j*), the contact-free area (*a*_*i*_ for cell *i*), and the *α*-parameter (half the ratio between the cell-cell surface tension *γ*_*cc*_ and cell-medium surface tension *γ*_*cb*_). At mechanical equilibrium, the *α*-parameter is inversely proportional to the angle between the surface of two adjacent cells *θ* (Young-Dupré equation). Blue, surface areas; red, surface tensions; magenta, angle. (**B,C**) Simulation of the compaction of the D_2d_ (B) and C_s_(2) (C) packing from *α* = 0.8 to *α* = 0.3 showing a topological transition to D_2d_ at *α* = 0.3 in (C). (**D**) Relative energy of the packings obtained computationally after compaction of the 13 rigid packings to *α* = 0.8, 0.55 and 0.3. Colour code corresponds to the topological proximity to C_s_(2) (magenta), D_2d_ (cyan) and other rigid packings (grey), and may differ from the initial packing used in the simulation due to topological transitions. (**E,F**) tSNE projection of the morphomap of minimally rigid packings during *in silico* compaction. Each rigid packing is represented with a sequence of points connected from *α* = 0.8 to *α* = 0.3 and coloured as a function of the compaction parameter (E) and the topological proximity to C_s_(2), D_2d_ or any other rigid packing (F, respectively magenta, cyan and grey). (**G**) 2D schematic explaining the process of importing 3D segmentations in Surface Evolver as 3D objects. The labels (top-left) are imported into Surface Evolver as clouds of points from which a convex shape is generated with correct contacts and positions (bottom-left). The first optimisation generates an uncompacted object with *α* = 0.95 ensuring cell volume consistency with the embryo (bottom-right). Further simulated compaction is then performed up to the relevant experimentally calculated *α*-parameter (top-right). See also Methods. (**H**) Visual comparison of the 3D segmentation of a representative mouse embryo (first row) and the corresponding *in silico* simulation (second row) at *α* = 0.7 (beginning of the simulation, left column) and *α* = 0.4 (end of the simulation, right column). Time after the beginning (white, top-left corner) and before the end (red, top-right corner) of the 8-cell stage. Scale bar, 25 *µ*m. (**I**) Distribution of the pair-wise distance in-between embryos (white, 10 embryos, 45 pairs) and between embryos and simulations (red, 10 embryos and 10 simulations) as a function of the compaction parameter from *α* = 0.65 (low compaction) to *α* = 0.4 (high compaction). (**J**) Schematic summary of the physical model. At very low compaction (*α* = 0.9), rigid packings are energetically equivalent, while compaction biases the relative energy of the different local minima, which favours noise-induced topological transitions (*α* = 0.6). At higher compaction rate (*α* = 0.3), multiple topologies have converged to D_2d_, although some topologies (D_3d_, C_2v_(2), C_s_(3) and C_s_(1a)) would require higher noise or longer time scales. See also Figure S4A,B.

To challenge its predictions further and more quantitatively, we compared embryo morphodynamics with their simulated counterparts, by using real embryo structure and geometry and topology at the start of the 8-cell stage as an initial condition to start the simulation (Figure 4G). The model could successfully recapitulate both the transitions from non-rigid to rigid packings as well as the geometrical convergence (Figure 4H). In particular, we found that the geometrical distance between embryos and their corresponding simulations is significantly smaller than the geometrical distance in random pairs of embryos (Figure 4I). Taken together, our *in-silico* model based on the surface energy minimisation was able to recapitulate the 8-cell embryo morphogenesis and predict geometrical and topological convergence to D_2d_. The attraction to D_2d_ can thus be explained by having the least surface energy for every tested *α*-parameter (Figures 4D and S4E), which suggests that with a sufficient amount of time and some level of dynamic changes and fluctuation in cell shape (Figure S4A,B), all packings would eventually converge to D_2d_ (Figure 4J).

### Compaction and surface contractility drive topological transitions

Our simulations predict that compaction plays a key role in the convergence towards a well-defined topological structure, by favouring a specific cellular packing as well as lowering barriers for transitions. Consistently, we found that the increase in probability towards the D_2d_ topology correlated strongly with the compaction parameter *α* (Figures 5A and S5A,B), increasing from 0.0% to 28.2% for *α* between 0.8 and 0.6 respectively, and from 27.6% to 70.0% for *α* < 0.45. Although the decrease of *α* varied significantly from embryo to embryo, the probability of D_2d_ topology correlated much more strongly with *α* than with time (Figure 3F), arguing that compaction is a primary driver of the convergence to D_2d_ during the course of the 8-cell stage. To functionally test this, we first generated mT embryos that lack the maternal allele of Myh9 (hereafter referred to as mMyh9^+/-^). As Myh9 is the specific isoform of myosin heavy chain that is required to generate surface tensions, in its absence embryos fail to compact properly and build cortical contractility (Maître et al., 2015) (Figure 5B,C and Video S5). In line with the predictions, mMyh9+/-embryos showed no topological transitions to D_2d_ for compaction parameter until as low as 0.55 (Figures 5D and S5C), while non-rigid packings were overly-represented throughout the 8-cell stage (mean ± s. d. = 75.6% ± 7.8) and embryos did not converge topologically nor geometrically (Figures 5E and S5D). Furthermore, using *para*-aminoblebbistatin (PAB, 10 *µ*m), a photostable myosin II inhibitor (Várkuti et al., 2016), we obtained embryos with a milder effect on compaction and showing intermediate topological and geometrical outcome (Figure S5E-I and Video S6). Altogether, these findings confirm the predictions of our model and demonstrate that compaction and surface contractility drive topological transitions and spatial convergence in the early mouse embryo.

**Figure 5.**
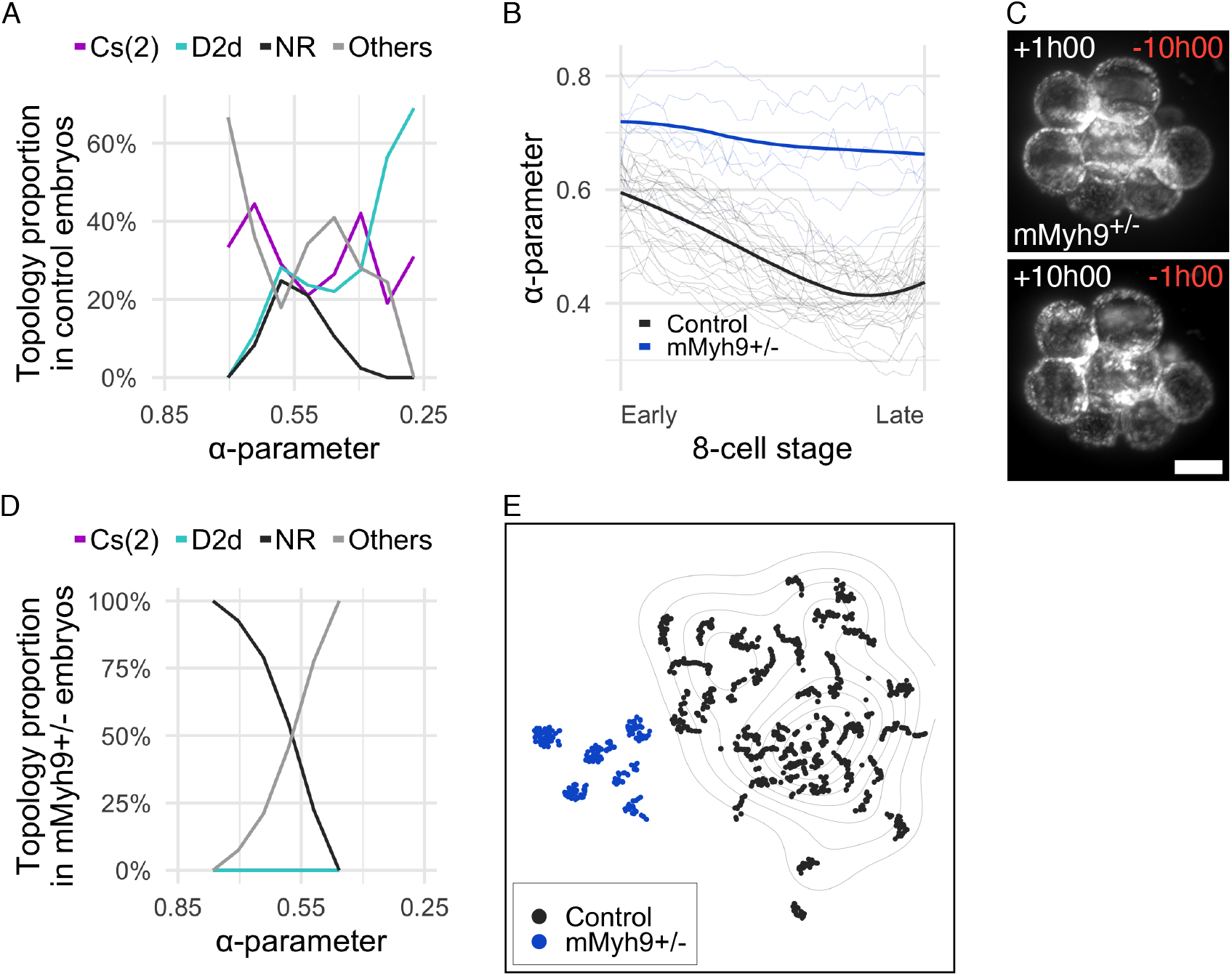
Compaction and surface contractility drive topological transitions. (**A,D**) Proportion of C_s_(2) (magenta), D_2d_ (cyan), other rigid packings (Others, grey) and non-rigid packings (NR, black) as a function of the compaction parameter in control mouse embryos (A, *n* = 29 embryos) and mMyh9^+/-^ mutants (D, *n* = 6 embryos). (**B**) Normalised time course through the 8-cell stage of the mean *α*-parameter for control mouse embryos (black, *n* = 29 embryos) and mMyh9^+/-^ mutants (blue, *n* = 6 embryos). Light colours, individual tracks. See also Figure S5D. (**C**) Max projection of a representative live imaging of mMyh9^+/-^ mutants at the beginning (top) and the end (bottom) of the 8-cell stage (*n* = 6 embryos). Time after the beginning (white, top-left corner) and before the end (red, top-right corner) of the 8-cell stage. Scale bar, 25 *µ*m. See also Video S5. (**E**) tSNE projection of the morphomap of control mouse embryos (black, *n* = 29 embryos) and mMyh9^+/-^ mutants (blue, *n* = 6 embryos). Isolines, density map at the end of the 8-cell stage of control embryos. See also Figure S5F.

### Variability in cleavage timing promotes robustness in morphogenesis and ICM-TE patterning

We thus far characterised the temporal variability in cleavage timing and the spatial convergence of the packings in the 8-cell embryo. Based on these characterisations, we next wished to investigate their impact on the forthcoming developmental stages, in particular on the first inside-outside cell fate patterning in the mouse embryo. First, we examined the impact of spatial convergence of the 8-cell embryo on the inside-outside cell segregation that happens in the 16-cell embryo. However, the number of potential rigid packings grows super-exponentially with the number of spheres involved, making the mapping of embryo packings yet unfeasible for the 16 cell stage (Holmes-Cerfon, 2016). To overcome this difficulty, we introduced a packing parameter that measures the deviation of outer cells from a certain distance to the embryo centre, inverted such that a higher value indicates a more packed embryo (Figure S6A and Methods). This packing parameter shows a high correlation with the 8-cell stage topological convergence (Figure 6A), justifying its use as a proxy in subsequent developmental stages. To determine the packing parameter at the 16-cell stage, we defined an outer cell as one belonging to the group with higher contact-free surface area, of the two groups forming a bi-modal distribution of the contact-free surface area (Figures 6B and S6B; Methods). When the packing parameter of the 16-cell embryos was computed using outer cells, it showed a clear correlation with that of the 8-cell embryos (Pearson correlation *R* = 0.658 (*P* < 0.01), Figure 6C) and the number of inner cells (Pearson correlation *R* = 0.591 (*P* < 0.01), Figure 6D), indicating that spatial convergence at the 8-cell stage facilitates the packing and hence the generation of inner cells in the 16-cell embryos. These findings show that spatial convergence and higher packing promote generation of inner cells in the early mouse embryo.

**Figure 6.**
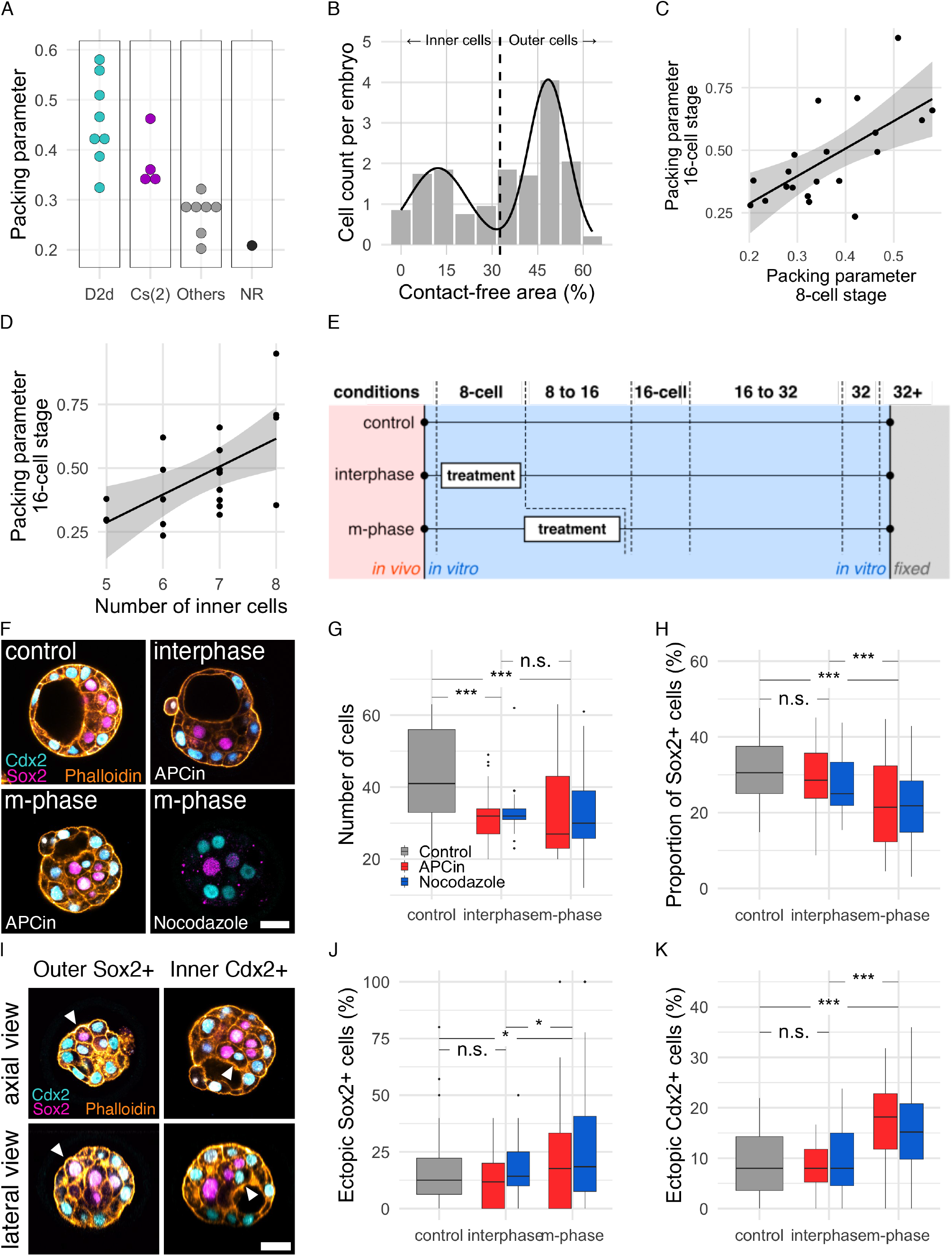
Variability in cleavage timing promotes robustness in morphogenesis and ICM-TE patterning. (**A**) Packing parameter at the 8-cell stage as a function of the topology of the embryo. *n* = 20 embryos. (**B**) Bi-modal count distribution of the contact-free area of individual cells (grey bars, *n* = 320 cells pooled from 20 embryos) fitted with the sum of an inner and an outer gaussian distribution (black line, mean ± s. d. = 12.1% ± 9.6 and 48.4% ± 6.5 respectively). Dashed line, cut-off between inner and outer cells, defined as the intersection point of the two gaussians (cut-off = 32.6%). (**C**) Packing parameter at the 16-cell stage as a function of the packing parameter at the 8-cell stage. Pearson correlation *R* = 0.658 (*P* < 0.01). Solid line, linear regression. Shaded ribbon, standard error. (**D**) Packing parameter as a function of the number of inner cells. Pearson correlation *R* = 0.591 (*P* < 0.01). Solid line, linear regression. Shaded ribbon, standard error. (**E**) Diagram of the synchronisation experiment describing the three conditions used in (F-H). Embryos were treated for 4 hours with APCin or Nocodazole at the beginning (group interphase, *n* = 25 embryos from 4 independent experiments and *n* = 25 embryos from 6 independent experiments for APCin and Nocodazole treatment respectively) or at the end (group m-phase, *n* = 43 embryos from 4 independent experiments and *n* = 72 embryos from 6 independent experiments for APCin and Nocodazole treatment respectively) of the 8-cell stage, or were not treated at all (group control, *n* = 57 embryos from 10 independent experiments). Embryos were fixed at the expected 32-cell stage (see Methods). (**F**) Representative cross-section of fixed embryos from the control group (top-left), the interphase group treated with APCin (top-right), the m-phase group treated with APCin (bottom-left) and the m-phase group treated with Nocodazole (bottom-right, stained without Phalloidin). Cyan, TE-fated cells (Cdx2); Magenta, ICM-fated cells (Sox2); Orange, F-actin (Phalloidin). Scale bar, 25 *µ*m. (**G,H,J,K**) Box plot of the number of cells (G), the proportion of Sox2+ cells (H), ectopic Sox2^+^ cells (J) and ectopic Cdx2^+^ cells (K) in embryos treated with APCin (red), Nocodazole (blue) or not treated (grey) in the three experimental groups as described in (E). Two-sided Mann-Whitney test; n.s., non-significant; *, p-value < 0.05; ***, p-value < 0.001. (**I**) Axial (top row) and lateral (bottom row) cross-section of fixed embryos from group m-phase, treated with APCin. Arrow head indicates ectopic Sox2^+^ (left column) and Cdx2^+^ (right column) cells. Colours and scale bar as in (F).

Next, we investigated the impact of variability in cleavage timing on the spatial convergence and precision in embryo patterning. To test the impact of temporal variability, we pharmaceutically synchronised mitotic entry at the 8-to 16-cell cleavage (Figure 6E) using an APC/C inhibitor (APCin) or a blocker of microtubule polymerisation (Nocadazole) during the 8-to 16-cell mitotic period (group “m-phase”), while control embryos undergo the same treatment during the inter-mitotic period, hence having no impact on division synchrony (group “interphase”). The APCin-treated embryos had a lower packing parameter (mean ± s. d. = 0.47 ± 0.18 and 0.37 ± 0.11 for untreated and APCin treated embryos respectively, Figure S6C-G), while maintaining a high number of inner cells (mean ± s. d. = 7.67 ± 1.03). Remarkably, this resulted in a lower proportion of the ICM-fated Sox2^+^ cells than in the control groups, both after APCin or Nocodazole treatment (Figure 6H). This indicates that synchronisation of the 4^th^ cleavage led to formation of the lower proportion of ICM-fated cells in the embryo. Moreover, treated embryos show significantly higher proportions of cells ectopically expressing Sox2 or Cdx2 (Figure 6I-K). Collectively, these results demonstrate that a certain degree of desynchronisation of the cleavages in the early mouse embryo enhances embryo packing, generation of the ICM fate, and precision in the inside-outside patterning.

## Discussion

In this study, we measured and manipulated spatial and temporal variabilities in early mammalian embryogenesis, which demonstrated that spatial and temporal variabilities are functionally linked and an asynchrony in cell divisions facilitates robust morphogenesis and patterning.

### Measurement and manipulation of temporal and spatial variabilities

Although cell-lineage has been established for many species, few studies focused on the cell-to-cell variability in spatial arrangement or cleavage timing (Anderson et al., 2017; Kelly et al., 1978; Olivier et al., 2010; Van et al., 1981; Villoutreix et al., 2016), with the exception of active desynchronisation described in Ascidian embryos (Dumollard et al., 2013; Dumollard et al., 2017; Sallé and Minc, 2022) and in human embryos (Mashiko et al., 2022; Roux, 1995). With a small number of cells and progressively accumulating variabilities in space and time, early mammalian embryos present an excellent opportunity to measure the building of variabilities during development.

To measure the spatial variability, we developed a new morphometric based on exponential-splines and compared embryos geometrically. While other methods have been used (Guignard et al., 2020; Kuang et al., 2022), including Fourier-shape descriptors (Agus et al., 2020; Ducroz et al., 2012; Tournemenne et al., 2014; Valizadeh and Babapour Mofrad, 2022) and dictionary-based shape-descriptors (Andrews et al., 2021; Saad et al., 2019; Tassy et al., 2006; Xiong and Sugioka, 2020), exponential splines encompass all geometrical hidden features (including volume, position, contacts, and compaction) with an arbitrary number of parameters and only a few assumptions, offering a generic tool to build developmental morphomaps. Regardless of the method, the geometrical data are ultimately reduced to fewer parameters which inherently approximate the actual cell shape. By smoothing the surface of the object, spline approximation of the surface minimise technical noise introduced by the imaging and processing techniques, such as voxel anisotropy or segmentation errors, while preserving the key geometrical features. Furthermore, topological analysis offers a theoretical framework which can be used to add landmarks to morphomaps, thus enhancing their interpretability in terms of packing. While it is possible to generate a morphomap and classify topologies for the 16-cell stage or later, the number of minimally rigid topologies increases super-exponentially with the number of cells (Holmes-Cerfon, 2016), so that further conceptual and technical advancements would be required to perform such exhaustive classification. The spatial variability at the beginning of an inter-mitotic period may be largely influenced by previous cell divisions, as cytokinesis generates force separating two daughter cells, thereby rearranging embryonic packings locally and abruptly, in a stochastic fashion. In particular, cleavage synchrony and axis of division could play a key role. Our present study focused on changing timing, and experimental synchronisation revealed a functional link between temporal and spatial variabilities in embryogenesis, however, future studies may directly disrupt the spatial arrangement of the cells.

It will also be interesting to examine whether temporal variability in cleavage timing is generated by stochastic and/or deterministic mechanisms (Froese, 1964; Sandler et al., 2015; Soltani et al., 2016). In addition, although we used cleavage timing to characterise the temporal variability in developmental progression, future studies may consider using, for example, the onset of certain gene expression as a timer of the development. Overall, since spatio-temporal variability is widespread, it will be interesting to explore its possible role in various developmental contexts, as cell fate, segregation and form are being established further.

### Optimal spatio-temporal variability facilitates robust morphogenesis and patterning

We showed that changes in cortical contractility drives not only compaction (Maître et al., 2016; Samarage et al., 2015), but also topological transitions by lowering the surface energy of specific packings, thus favouring embryo convergence towards an optimally packed shape. There is, however, an infinite number of possible trajectories between two defined topologies. What makes the embryo follow one trajectory over another is yet to be elucidated and depends on the shape of the energy landscape, passive minimisation of surface energy and active cellular mechanisms. Although additional mechanisms may constrain embryos to a limited subset of the morphomap, e.g., a geometrical constraint of the zona pellucida, as shown with the eggshell in *C. elegans* (Seirin-Lee et al., 2022), or the possible spatial coupling between sister cells at the 8-cell stage (Lim et al., 2020), we showed that minimisation of contractility-driven surface energy plays a critical role and are sufficient to explain key features of geometrical and topological convergence.

In this study, we discussed three different, yet interdependent, expression of variabilities: cleavage asynchrony, cell shape fluctuation and packing heterogeneity. The first two may be described as temporally varying input mechanisms which impact the latter spatial variability. Cell contractility is thought to generate stochastic fluctuations of cellular shape and to facilitate the loss and the formation of cell-cell contacts. Some minimal level of fluctuations is therefore required for topological transition. In synchronous cleavages, the cell shape changes are dominated by the impact of cell divisions, which results in dramatic geometrical and topological changes leading to disrupted morphogenesis and patterning. Interestingly, the idea that noise is not detrimental to reach a given stable state, but instead might be necessary to avoid being trapped in local minima, has been heavily explored in computer science as well as in physics and chemistry (Horsthemke and Lefever, 2006; van Kampen, 2007), as exemplified for instance in the method of stochastic gradient descent for finding optima in systems with complex energy landscapes (Arbib, 1998; Tsypkin, 1971). We thus conjecture that intermediate levels of variability – spatial or temporal, either arising from cleavages or contractility fluctuations – might be optimal for converging towards stereotypical embryo shapes and pattern.

### Shared and distinct features across mammals highlight essential processes ensuring developmental robustness

Previous work showed that mouse, rabbit, monkey and other species exhibit unique and shared developmental features in gene expression, epigenetic regulation, or as a whole with interspecies chimeras (Simerly et al., 2011; Okamoto et al., 2011; Wu et al., 2016; Nakamura et al., 2016; Fu et al., 2020; Bouchereau et al., 2022). However, little was known about cellular morphogenesis, cleavages, and their variabilities. Here we demonstrated that the cellular morphogenesis of mouse, rabbit and monkey embryos can be described in a shared morphomap, revealing key similarities and differences between species. Rabbit embryos showed a much higher spatial variability and a low proportion of D_2d_ packing, which is in line with our model derived from mouse embryos given that little compaction is observed at the 8-cell stage. Instead, the compaction in rabbit embryos starts when embryos reach the 32- and 64-cell stage (Koyama et al., 1994), which opens the possibility for a late topological and geometrical convergence. By contrast, monkey embryos display robust geometrical and topological patterns despite the absence of compaction at the 8-cell stage. This suggests that cleavage asynchrony may facilitate robust morphogenesis, by promoting an optimal packing from the beginning of the 8-cell stage. Alternatively, the zona pellucida appears to expert higher spatial constraints to monkey embryos than to mouse embryos (see Figures 2E and S2C, in comparison to Figures 2A and S2A), and this may also contribute to the efficient convergence and packing (Giammona and Campàs, 2021; Seirin-Lee et al., 2022).

Overall, the dynamics of the variability in cleavage timing is closely linked to the spatial organisation of the cells within the embryo, and ultimately to morphogenesis and patterning. Our finding of species-specific dynamics of variability in cleavage timing suggests an intriguing possibility that the temporal variability may be an evolutionary trait generating an optimal spatial noise, ultimately ensuring robustness in embryogenesis.

## Acknowledgments

We are grateful to the members of the Hiiragi laboratory for discussions and comments on the manuscript; Ramona Bloehs, Stefanie Friese, Wibke Schwarzer and Lidia Pérez for their technical support; Vera Janssen for establishing the PAB protocol; members of the Tsukiyama group for the animal care with monkeys, in particular Hideaki Tsuchiya and Masataka Nakaya; Unité Commune d’Expérimentation Animale (UCEA) for the animal care with rabbits; the EMBL electronic and mechanical workshops and the EMBL animal facility for their support; We thank Luxendo for the close collaboration in developing the light-sheet microscopy for mammalians; D.F. was supported by fellowships from the EMBL Interdisciplinary Postdoc Program (EIPOD) under Marie Sklodowska Curie Actions (COFUND III RTD) and T.I. by JSPS Overseas Research Fellowship. We thank the generous support to B.C-M of the *field of excellence* “Complexity of life in basic research and innovation” of the University of Graz. The Hiiragi laboratory was supported by the EMBL, and currently by the Hubrecht Institute, the European Research Council (ERC Advanced Grant “SelforganisingEmbryo”, grant agreement 742732), Stichting LSH-TKI (LSHM21020) and JSPS KAKENHI grant number JP21H05038.

## Author contributions

Conceptualization, D.F., B.C.-M., E.H., T.H.; Methodology, D.F., B.C.-M., A.Kr., V.U., E.H., T.H.; Software Programming, D.F., B.C.-M., A.W., V.U.; Validation, D.F., B.C.-M., E.H., T.H.; Formal Analysis, D.F.; Investigation, D.F., P.M., A.Ki., T.I., C.I., N.D.; Resources, T.T., A.Kr., V.D., T.H.; Data Curation, D.F., A.Ki, J.G., A.S.; Writing – Original Draft, D.F., B.C.-M., E.H., T.H.; Writing – Review & Editing, D.F., B.C.-M., A.Ki, V.U., E.H., T.H; Visualization, D.F., B.C.-M.; Supervision, D.F., B.C.-M., T.T., A.Kr., V.D., V.U., E.H., T.H.; Project Administration, D.F., T.H.; Funding Acquisition, D.F., T.H.

## Materials and Methods

### Mouse work

We performed mouse animal work in the Laboratory Animal Resources (LAR) at the European Molecular Biology Laboratory with permission from the Institutional Animal Care and Use Committee (IACUC) overseeing the operation (IACUC number TH11 00 11). LAR is operated as stated in international animal welfare rules (Federation for Laboratory Animal Science Associations guidelines and recommendations). Mouse colonies are maintained in specific pathogen-free conditions with 12– 12 h light-dark cycle. All mice used for experiments were older than 8 weeks.

### Rabbit work

We performed rabbit animal work following the International Guidelines on Biomedical Research involving animals, as promulgated by the Society for the Study of Reproduction, and with the European Convention on Animal Experimentation. The researchers involved in work with the rabbits were all licensed for animal experimentation by the French veterinary services. The rabbit experimental design was carried out under the approval of national ethic committee (APAFIS #2180-2015112615371038v2) and under the approval of the local ethic committee (Comethea n°45, registered under n°12/107 and n°15/59).

### Monkey work

We performed monkey animal work with female cynomolgus monkeys (*Macaca fascicularis*), of ages ranging from 6 to 11 years. The light-dark cycle was maintained as 12-12 hours of artificial lighting from 8 a.m. to 8 p.m. Each animal was fed 20 g/kg of body weight of commercial pellet monkey chow (CMK-1; CLEA Japan) in the morning, supplemented with 20–50 g of sweet potato in the afternoon. Water was available *ad libitum*. Temperature and humidity in the animal rooms were maintained at 25 ± 2°C and 50 ± 5%, respectively. The animal experiments were appropriately performed by following the Animal Research: Reporting in Vivo Experiments (ARRIVE) guidelines developed by the National Centre for the Replacement, Refinement & Reduction of Animals in Research (NC3Rs), and also by following “The Act on Welfare and Management of Animals” from Ministry of the Environment, “Fundamental Guidelines for Proper Conduct of Animal Experiment and Related Activities in Academic Research Institutions” under the jurisdiction of the Ministry of Education, Culture, Sports, Science and Technology, and “Guidelines for Proper Conduct of Animal Experiments” from Science Council of Japan. All animal experimental procedures were approved by the Animal Care and Use Committee of Shiga University of Medical Science (approval number: 2021-10-4).

### Mouse lines

The following mouse lines were used in this study: (C57BL/6xC3H) F1 for WT, mTmG (*Gt(ROSA) 26Sor*^*tm4(ACTB-tdTomato,-EGFP)Luo*^) (Muzumdar et al., 2007), CAG H2B-EGFP (*Tg(HIST1H2BB/EGFP)1Pa*) (Hadjantonakis and Papaioannou, 2004) and *Myh9*^*tm5RSad*^ (Jacobelli et al., 2010). Double transgenic mTmG ; H2B-EGFP line was generated by breeding mTmG (homozygous or heterozygous) females with H2B-EGFP males. Transgenic mMyh9 ; mTmG ; H2B-EGFP embryos were generated by maternal deletion of Myosin-9 using ZP3-Cre (*Tg(Zp3-cre)93Knw*) mice (de Vries et al., 2000) and *Myh9*^*tm5RSad*^ mice. Myh9^tm5RSad/tm5RSad^ ; Zp3^Cre/+^ mothers were bred with mTmG ; H2B-EGFP fathers. mG line was generated by maternal excision of the mT sequence using ZP3-Cre mice and mTmG mice. Genotyping was done using the following primers:

- mTmG and mG lines: 250 b.p. (transgenic) and 330 b.p. (wild-type) PCR product size with primers oIMR7318 (CTCTGCTGCCTCCTGGCTTCT), oIMR7319 (CGAGGCGGATCACAAGCAATA) and oIMR7320 (TCAATGGGCGGGGGTCGTT),
- CAG H2B-EGFP: 900 b.p. (transgenic) PCR product size with primers CAG-F (GGCTTCTGGCGTGTGACCGGC) and EXFP-R (GTCTTGTAGTTGCCGTCGTC); 324 b.p. (wild-type) PCR product size with primers oIMR7338 (CTAGGCCACAGAATTGAAAGATCT) and oIMR7339 (GTAGGTGGAAATTCTAGCATCATCC); PCR must be done separately,
- *Myh9*^*tm5RSad*^: 770 b.p. (transgenic) and 600 b.p. (wild-type) PCR product size with primers Myh 9F (ATGGGCAGGTTCTTATAAGG) and Myh 9R (GGGACACAGTGGAATCCCTT),
- ZP3-Cre: 300 b.p. (transgenic) PCR product size with primers *cre upper* (TGCTGTTTCACTGGTTGTGCGGCG) and *cre lower* (TGCCTTCTCTACACCTGCGGTGCT).

### Recovery of mouse embryos

Embryos were recovered from super-ovulated female mice. For superovulation, intraperitoneal injection of 5 international units (IU) of pregnant mare’s serum gonadotropin (PMSG, Intervet, Intergonan) and following injection of 5-IU human chorionic gonadotropin (hCG; Intervet, Ovogest 1500) 48 h later were performed. 4-cell stage embryos were recovered at E2.0 by inserting a needle in the ampulla and flushing the oviduct and the uterus with KSOMaa including HEPES (H-KSOMaa; Zenith Biotech, ZEHP-050). Recovered embryos were washed three times in drops of KSOMaa (Zenith Biotech, ZEKS-050) and then transferred into 10 µl drops of KSOMaa covered with mineral oil (Sigma-Aldrich, M8410). Embryos were maintained in an incubator (Thermo Fisher Scientific) at 37ºC with 5% CO2.

### Recovery of rabbit embryos

New Zealand White female rabbits (20–22 weeks old) were super-ovulated using 5 subcutaneous administrations of porcine follicle stimulating hormone (pFSH, Merial, Stimufol) for 3 days before mating: two doses of 5 µg on day 1 at 12 hours intervals, two doses of 10 µg on day 2 at 12 hours intervals, and one dose of 5 µg on day 3 followed 12 hours later by an intravenous administration of 30IU hCG (Intervet, Chorulon) at the time of mating (natural mating). Embryos were collected from oviducts and perfused with DPBS (Thermo Fisher Scientific, 14190) at 19 hours post-coitum (h.p.c.) to obtain embryos at 2 pronuclei stage. Embryos were maintained in 0.5mL of TCM199-HEPES (Biochrom, F0665) + 10% foetal bovine serum (FBS, Thermo Fisher Scientific, 10500) + 0.5% of penicillin/streptomycin (Thermo Fisher Scientific, 15140) in a 4-well dish (Nunc, 176740) until the microinjection.

### Intracytoplasmic sperm injection (ICSI)

Monkey oocyte collection was performed as described previously (Tsukiyama et al., 2019). Briefly, two weeks after the subcutaneous injection of 0.9 mg of a gonadotropin-releasing hormone antagonist (Takeda Chemical Industries, Leuplin for Injection Kit), a micro-infusion pump (ALZET Osmotic Pumps, iPRECIO SMP-200) with 15 IU/kg human FSH (hFSH, Merck Biopharma, Gonal-f) was embedded subcutaneously under anaesthesia and injected 7 µL/h for 10 days. After the hFSH treatment, 400 IU/kg human chorionic gonadotropin (hCG, Asuka Pharmaceutical, Gonatropin) was injected intramuscularly. Forty hours after the hCG treatment, oocytes were collected by follicular aspiration using a laparoscope (Machida Endoscope, LA-6500). Cumulus-oocyte complexes (COCs) were recovered in alpha modification of Eagle’s medium (MP Biomedicals, 09103112-CF), containing 10% serum substitute supplement (Irvine Scientific, 99193). The COCs were stripped off cumulus cells with 0.5 mg/ml hyaluronidase (Sigma-Aldrich, H4272). ICSI was carried out on metaphase II (MII)-stage oocytes in mTALP+HEPES (Parrish et al., 1988) with a micromanipulator. Fresh sperm were collected by electric stimulation of the penis with no anaesthesia.

### *In vitro* transcription

H2B-EGFP mRNA was transcribed from plasmid #1 pCS-H2B-EGFP (Megason, 2009) (Addgene, Plasmid #53744). To construct pCS2-myrTagRFP-T (plasmid #2), TagRFP-T fused with eight amino acids, GSSKSKPK, at the N-terminus for myristoylation was amplified (primer set: 5’-GGATCCATGG GCAGCAGCAA GAGCAAGCCC AAGAGCGAGC TGATTAAG-3’ and 5’-CTCGAGTCAC TTGTGCCC-3’) and cloned into the BamHI-XhoI sites of pCS2+. To construct pcDNA3.1-membrane-tdTomato-T2A-H2B-GFP-poly(A83) (plasmid #3), an amplified PCR product from pCAG-TAG (Addgene, Plasmid #26771) was cloned into the KpnI-NotI sites of pcDNA3.1-H2B mCherry-poly(A83) (Yamagata et al., 2005). The vector #3 was linearized with XhoI and treated with 0.5% SDS, 0.2 mg/mL Proteinase K (Thermo-Fisher, QS0510) for 30 min at 50°C, purified with phenol-chloroform, and precipitated with ethanol.

Then, the purified vectors were used as a template for *in vitro* transcription. The mRNA from plasmid #1 and #2 was transcribed using the mMESSAGE mMACHINE SP6 Transcription Kit (Thermo Fisher Scientific, AM1340) and purified with the NucleoSpin RNA Clean-up XS kit (Macherey-Nagel, 740902). The mRNA from plasmid #3 was transcribed using the mMESSAGE mMACHINE T7 Transcription Kit (Thermo Fisher Scientific, AM1344) and purified with the MEGAclear Transcription Clean-Up Kit (Thermo Fisher Scientific, AM1908).

### Micro-injection of mRNAs

Rabbit zygotes were micro-injected into the cytoplasm with a mRNA’s mix of H2b-GFP (50 ng/µL final concentration) and myrTagRFP-T (150 ng/µL final concentration) in water. Microinjections were performed using a DIC inverted microscope (Olympus, IX71) equipped with micromanipulators (Eppendorf, TransferMan NK) and electronic microinjector (Eppendorf, femtojet injector). After microinjection, unlysed embryos were cultured during 2 to 3 hours in 40 µL micro drops of TCM199 (Sigma-Aldrich, M4530) + 10% foetal bovine serum (FBS, Thermo Fisher Scientific, 10500) + 0.5% penicillin/streptomycin (Thermo Fisher Scientific, 15140) under mineral oil (Sigma-Aldrich, M8410) at 38.5°C under 5% CO2.

For microinjection in monkey eggs, the mRNA was co-injected with sperm during ICSI. The sperm were washed in a drop of 300 ng/µL mRNA, and co-injected into the MII-stage oocytes. Following co-injection, embryos were cultured in monkey culture medium (CMRL 1066 Medium (Thermo Fisher Scientific, 21540026) supplemented with 20% FBS) at 38°C in 5% CO2 and 5% O2.

### Transport of microinjected rabbit zygotes

Two to three hours after microinjection, 0.5 mL tubes were over-filled with M2 (Sigma-Aldrich, M7167) at 38.5°C to remove air bubbles. Tubes were placed into an insulated box (Polystyrene foam transport box) with the Velvet cooling elements (Velvet, SVE2) and antifreeze elements (Velvet, AF1) to maintain 8°C during 36 to 48 hours. The samples were sent from the collection site (Jouy-en-Josas, France) to the European Molecular Biology Laboratory (Heidelberg, Germany) via a regular courier service overnight. Before reception of the embryos, rabbit culture medium (TCM199 (Sigma, M4530) + 10% FBS (Thermo Fisher Scientific, 10500) + 0.5% penicillin/streptomycin (Thermo Fisher Scientific, 15140)) was freshly prepared and stored at 38.5ºC and 5% CO2 for at least an hour. Upon reception of the embryos, the temperature inside the box was checked (8ºC), the embryos transferred to a 4-well dish (Nunc, 176740) with M2 (Sigma, M7167) at room temperature for 5 minutes, washed four times in 40 µL drops of equilibrated and pre-warmed rabbit culture medium, then transferred to 10 µL drops of equilibrated and pre-warmed rabbit culture medium covered with mineral oil (Sigma, M8410). Embryos were maintained in an incubator (Thermo Fisher Scientific) at 38.5ºC with 5% CO2 for one hour until the beginning of the imaging.

### Transport of microinjected monkey embryos

Forty-five hours after microinjection, 4-cell stage monkey embryos were transferred into 300 µL monkey culture medium (CMRL 1066 Medium supplemented with 20% FBS) covered with 1 mL mineral oil in a 1.5 mL tube. Tubes were placed into 38.5ºC water in a vacuum bottle in an insulated box to maintain temperature during 1.5 hours of transportation from collection site (Shiga University of Medical Science, Japan) to the imaging site (Kyoto University, Japan). Upon arrival, embryos were washed three times with pre-warmed monkey culture medium, then transferred to 10 µL drops of equilibrated and pre-warmed monkey culture medium covered with mineral oil. Embryos were maintained in an incubator (PHCbi, MCO-170MUV) at 38°C with 5% CO2 and 5% O2 for one hour until the beginning of the imaging.

### Chemical treatments

For Myosin inhibition (Figure 5), 4-cell stage embryos were washed and cultured in pre-equilibrated KSOMaa (Zenith Biotech, ZEKS-050) supplemented with DMSO (vehicle; Sigma-Aldrich, D2650) and *para*-aminoblebbistatin (10 *µ*m; Optopharma, DR-Am-89) for 24 hours until the 16-cell stage. For the synchronisation of cell mitoses at the 4^th^ cleavage (Figure 6), 8-cell stage embryos were washed and cultured at the appropriate timing for 4.5 hours in pre-equilibrated KSOMaa supplemented with 1:1000 DMSO (control group), APCin (100 *µ*m; Tocris Bioscience, 5747) or Nocodazole (0.5 *µ*m; Sigma-Aldrich, M1404). Embryos were then washed in pre-equilibrated KSOMaa. The interphase group was formed using early uncompacted 8-cell stage embryos, while the m-phase group was formed using late compacted 8-cell stage embryos. Embryonic stage (early or late 8-cell stage) was determined by visual inspection with bright-field binocular microscope (Zeiss, Discovery.V8).

### Generation of heterochronic embryos

mTmG and mG mice were super-ovulated four hours apart. Embryos were recovered at the 4-to 8-cell stage and imaged every 30 minutes in the InVi-SPIM to identify the beginning of the 8-cell stage. Pairs of one mT and one mG expressing embryos with 3 to 6 hours of difference were selected to generate heterochronic chimeras. First, one of the two embryos was randomly chosen to be the donor. To dissociate the donor embryo into single blastomeres, the zona pellucida (ZP) was removed by incubating the embryos in pronase (0.5% w/v Proteinase K in H-KSOMaa supplemented with 0.5% PVP-40 (Sigma-Aldrich, P0930)) covered with mineral oil for 2-3 minutes. Embryos were then washed 5 times in 10 µL drops of H-KSOMaa. Afterwards, the embryos were placed into a 50 µL drop of dissociation medium (Biggers et al., 2000) (KSOMaa without Ca^2+^ and Mg^2+^). Blastomeres were then dissociated in the drop by pipetting up and down in a narrow glass capillary (Brand, 708744). Dissociated blastomeres were incubated in KSOMaa drops covered with mineral oil until the host embryo was ready for the graft. To prepare the host, we used a micro-manipulation device (Narishige, MON202-D) mounted on a Zeiss Axio Observer Z1 microscope to create a slit in the ZP. First, the host embryos were placed in a 50 µL drop of H-KSOMaa covered with mineral oil in a glass-bottom dish (MatTek, P50G-1.5-14-F) and mounted on the microscope with temperature of the incubation chamber maintained at 37*°*C. To make a slit in the ZP of the host, a holding pipette and a pulled glass needle mounted on the micromanipulator were used. Both the holding pipette and pulled needle were custom-made from glass capillaries (Warner Instrument, GC100T-15) using a micropipette puller (Sutter Instrument, P-1000) and a microforge (Narishige, MF-900). The host embryo was maintained in place with the holding pipette and the glass needle was inserted tangentially to the embryo under the ZP and pulled in a perpendicular direction without damaging the cells, thus generating a slit in the ZP. Next, one of the donor’s blastomeres was randomly picked and inserted in the slit under the host’s ZP using another custom-made glass pipette mounted on the micromanipulator. The resulting 9-cell heterochronic chimeras were then transferred to KSOMaa and put back in the InVi-SPIM and imaged every half an hour until the 32-cell stage. Finally, timing of division was manually inspected from 8-to 16-cell stage and from 16-to 32-cell stage. Host and donor cells were identified based on the expression of mT or mG.

### Immunostaining

Embryos were fixed in 100µL of 4% paraformaldehyde (PFA, Electron Microscopy Sciences, 19208) in DPBS for 15 min at room temperature, washed three times for 5 minutes in DPBS with 0.1% Tween-20 (wDPBS, Sigma-Aldrich, P7949) + 1% bovine serum albumin (BSA, Sigma-Aldrich, 9647), permeabilised 15 minutes at room temperature in DPBS with 0.5% TritonX-100 (Sigma-Aldrich, T8787), washed three times for 5 minutes in wDPBS+1% BSA, and blocked 4 hours at room temperature in wDPBS+3% BSA. Embryos were then transferred into 70 µL of primary antibody solution in wDPBS+3% BSA and incubated overnight at 4ºC. Primary antibodies against Cdx2 (Biogenex, MU392A-UC) and Sox2 (Cell Signalling, D9B8N) were diluted at 1:150. Embryos were then washed three times for 5 minutes in wDPSB+1% BSA and transferred into 70 µL of secondary antibody solution in wDPSB+1% BSA for 2 hours at room temperature. Secondary antibodies conjugated with Cy5 against mouse Ig (Jackson ImmunoResearch, 715-175-150) and Alexa Fluor 488 Plus against rabbit Ig (Thermo Fisher Scientific, A11008) were diluted at 1:200. For embryos fixed after treatment with APCin (Figure 6), Phalloidin-Rhodamin (Thermo Fisher Scientific, R415) was added in the secondary antibody solution diluted at 1:200. Before imaging, embryos were washed three times for 5 minutes in wDPBS+1% BSA, transferred 10 minutes at room temperature in wDPBS+1% BSA supplemented with DAPI (1:1000, Thermo Fisher Scientific, D3751) to stain DNA, washed three times for 5 minutes in wDPBS+1%BSA and mounted in 5 µL drops of wDPBS for imaging the same day.

### Confocal microscopy

Imaging of immunostained embryos was performed with LSM 780 (Zeiss). C-Apochromat 403 1.1 NA water objective (Zeiss) was used.

### Light-sheet microscopy

Embryos were live-imaged using InVi-SPIM (Luxendo). Embryos were aligned in a V-shaped sample holder covered with transparent fluorinated ethylene propylene foil (FEP, Luxendo), in approximately 100 µL of culture medium covered with 200 µL of mineral oil to prevent evaporation. The sample holder was enclosed in an environmentally controlled incubation box with 5% CO2 and 5% O2 at 37ºC (mouse), 38ºC (monkey) or 38.5ºC (rabbit). For Figure 1, embryos were isolated in wells to facilitate their identification after imaging, fixation and immunostaining. To shape the wells, we used a flamed glass capillary (Marienfeld Superior, 2930210) with a spherical tip of approximatively 200 *µ*m of diameter to carefully press and stretch the FEP foil shaping up to 12 wells per sample holder. InVi-SPIM was equipped with a Nikon 25x/1.1NA water immersion detective objective and a Nikon 10x/0.3 NA water immersion illumination objective. The illumination plane and focal plane were aligned before each imaging session and maintained during the imaging. Images were taken by a CMOS camera (Hamamatsu, ORCA Flash4.0 V2) using software LuxControl (Luxendo). The lasers and filters used were 488 nm with BP525/50 and 561 nm with LP561 to image GFP and tdTomato/RFP fluorophores respectively. Exposure time for each plane was set to 30 ms. Imaging was done using a lateral resolution of 0.104 *µ*m/px or 0.208 *µ*m/px binned to 0.416 *µ*m for analysis. Due to technical difficulties with the motorised stage, the InVi-SPIM used a larger than requested step size between optical slices. To determine the real step size, we computed for each embryo the width:depth ratio and the height:depth ratio in voxels of all cells at every timepoint of the segmented 8-cell stage. Assuming that these ratio equals 1 on average, as with the width:height ratio, we determined the real step size in *µ*m for each embryo individually (Table S1). Imaging settings, including duration of imaging and time interval for each dataset, are summarised in Table S1.

### Generation of cluster of spheres *in-silico*

We describe the method to generate *in-silico* surrogates of packings such that they can be manipulated using the software Surface Evolver (Brakke, 1992), a classical tool to study the surfaces and topological transitions shaped by surface tension. Input data provides the geolocation of the centre of masses of cells, area of contact between cells and volume of each cell. Output data is a file compatible with Surface Evolver (file extension *fe*). For each cell, we created a cloud of points at distance *R* of its centre of mass, such that the enclosed volume corresponded to volume of the cell. For adjacent cells, we created a triangle – a simplex – that was shared among the two cells in contact. Then, we created the convex hull of the cloud of points, including the contact simplices if present, even though they can break locally the convexity of the object. This resulted in a structure made of triangular simplices to which we provided a consistent orientation such that they can be properly interpreted by Surface Evolver. Volumes were also given consistently with the volumes in the input data. The first optimization steps of Surface Evolver are sufficient to generate a structure akin to the input data, both at the topological and geometrical level, and also to remove the potential non-convex regions generated at the contact points. The surface tensions can then be modified such that one reproduces the compaction process. See also Figure 4G for a schematic description of the process.

### *In-silico* compaction and energy minimisation

Following previous findings that cell-cell adhesion during mouse compaction could be described well by the analogy to soap bubble (Maître et al., 2015; Maître et al., 2016), the energy of a given configuration is defined by

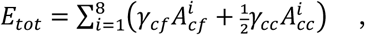

where we have summed by all eight cells their cell-fluid energy (*γ*_*cf*_ being the cell-fluid tension and 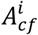 the cell-fluid area of cell *i*) and their cell-cell energy (*γ*_*cc*_ being the cell-cell tension and 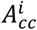 the area of cell *i* in contact with other cells, note that the factor ½ is to avoid double summation). We set all cellular volumes to unity. We can set *γ*_cc_ = 1 without loss of generality, so that the minimization problem depends only on the tension ratio *α* = *γ*_*cc*_/*γ*_*cf*_. It has been shown previously that this parameter *α* can be estimated from the angles between cell-cell contacts and the fluid interface (Young-Dupré equation, see also Figure S2E and 4A). Note that 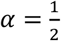 corresponds to soap bubbles (all tensions are equal and all angles are 120º), *α* = 0 corresponds to only cell-fluid tension so that the embryo will be perfectly spherical regardless of the contact topology, and *α* > 1 corresponds to the non-adhesive case as it becomes energetically favourable for cells to dissociate (as cell-cell tensions are larger than cell-fluid tensions).

To explore how each of the 13 rigid packings evolve with compaction, we first initialized the software with the 13 possible packings and a very small adhesion strength (*α* = 0.95), which ensured that cells showed very small cell-cell contacts, and no tricellular junctions, in analogy with adhesive hard colloids (Arkus et al., 2009). We then slowly decremented *α* by 0.05, each time allowing for equilibration to a minimal energy configuration. This allowed us to calculate how energies of each configuration evolve as a function of *α*. Importantly, while for *α* close to 1, energies of all 13 packings were very similar, as expected theoretically, decreasing *α* broke the degeneracy of the system. We found that D_2d_ was always the lowest energy configuration, followed by the C_s_(2) packing (Figure 4D). The local stability of these energy minima was verified in Surface Evolver by coarsening and remeshing the surface, effectively adding noise and checking whether the system always converged towards the same energy configuration. Interestingly, we found that 2 of the 13 packings (C_s_(5) and C_2v_(1)) disappeared upon compaction, as the associated deformations create new contacts which make the system converge to D_2d_ even in the absence of noise. For the other packings, convergence towards the global minimum (D_2d_) required T1 topological transitions, which is associated to energy costs in foam models. Indeed, we found that the introduction of a large amount of remeshing noise in the system was necessary to allow these packings to converge towards D_2d_ via topological transitions (Figure 4E,F and S4A,B). Note that two of the most anisotropic packings (C_2v_(2) and D_3d_) could not converge towards D_2d_, at least within the noise values possible with the Evolver package. It is possible that these highly anisotropic packings are unfavoured by other factors, such as asynchronous divisions.

### Nearest neighbour algorithm

For cell tracking, registration or alignment of two sets of 3D points (P_A_ and P_B_) or 3D segmentation (S_A_ and S_B_), we developed a nearest neighbour algorithm in Python, available in the package *morphomap* (see section *Code availability*). For segmentation, the centre of mass of each cell in S_A_ and S_B_ was used to generate P_A_ and P_B_ respectively. First, we computed C_A_ and C_B_, the centres of mass of P_A_ and P_B_ respectively. Second, P_A_ and P_B_ were translated to the origin such that P_A0_ = P_A_-C_A_ and P_B0_ = P_B_-C_B_. Third, we generated 125,000 transformation matrices including rotation (10 values between 0º and 360º for pitch, roll and yaw) and scaling (5 values 0.5, 0.75, 1.0, 1.25 and 1.50 along the X, Y and Z axis), corresponding to 125,000 transformations tP_B0_ of P_B0_. Pair-wise Euclidean distances between points of tP_B0_ and P_A0_ were computed and sorted. The pair were assigned once, in order from the smallest distance to the biggest distance. If tP_B0_ had unassigned cells after the first round, a second round of assignation was done, assuming mitosis. The alignment score was computed as the sum of the squared distance between assigned points of P_A0_ and tP_B0_. The best three scores were kept and a new set of 125,000 transformation matrices was generated exploring the grid cell around the selected values, progressively refining a grid search of rotations and scaling. The algorithm stopped after 3 iterations (corresponding to variations of 0.36º for the rotation and 0.025% for the scaling). In case of segmentation, the score was weighted by the Jaccard index between segmented cells (number of voxels in common divided by the total number of voxels). The nearest neighbour algorithm produced a translation vector, rotation parameters, scaling parameters and a list of cell assignment. Computation was distributed on the EMBL’s cluster.

### Tracking and manual curation

For mouse embryos, nuclear centres were detected using a Difference of Gaussians algorithm (Faure et al., 2016) (DoG). The best parameters for the DoG algorithm were found by a grid-search procedure which explored thousands of different configuration parameters simultaneously using EMBL’s computing cluster. The best output was manually chosen and used for tracking using our nearest neighbour algorithm. Manual curation from the 4-cell stage to the 32-or 64-cell stage was performed by one operator using the software Mov-IT (Faure et al., 2016) and validated by a second operator. All the cells were inspected. Alternatively, for rabbit and monkey embryos, we used a custom-made cell detection and tracking algorithm (unpublished) based on nuclear segmentation with Cellpose (Pachitariu and Stringer, 2022; Stringer et al., 2021) and semi-automatic shape tracking using Napari (Sofroniew, Nicholas et al., 2022).

### Identification of cell fate and back-tracking

Identification of inner-cell-mass (ICM) and trophectoderm (TE) progenitor cells was done by fixation and immunostaining of embryos less than 15 minutes after the end of the live imaging. Sox2 and Cdx2 positive cells (respectively ICM- and TE-fated cells) were identified by the signal intensity and mapped on the last timepoint of the live-imaged dataset. The alignment of the cells from the confocal imaging of immunostained embryos and the cells from the corresponding live-imaged embryos was done using our nearest neighbour algorithm. The identity of cells as determined by the immunostaining was then transferred to the last time point of the live-imaged dataset. Finally, cell identity was propagated backwards with a higher priority to TE-fated cells.

### Measurement of cleavage timing variability

Cells were annotated with their generation number determined by the number of cells at the beginning of the imaging (zygote stage being the 1^st^ generation). After each mitosis, daughter cells’ generation was incremented. The variability in division timing at the *n*^th^ cleavage was measured as the standard deviation of the timing of division (in hours) of the cells of the *n*^th^ generation in one embryo. When multiple embryos were pooled, the timing of division at one generation was first centred to zero for each embryo before computing the standard deviation of all the pooled cells at that generation. Embryos with less than 80% of cells dividing to the *n*^th^ generation during the imaging period were removed from the analysis.

### Time and spatial difference between cells

For each embryo, all unique pair of cells of the same generation were generated. The time difference was measured as the time (in hour) between the divisions of the cells in the pair. The spatial difference was measured as the Euclidean distance between the two cells 15 minutes before the first mitosis.

### Membrane segmentation and curation

The segmentation pipeline used to process the 3D images of the membrane signal uses the PlantSeg package (Wolny et al., 2020) based on previous work done in electron microscopy images of neural tissue (Beier et al., 2017; Funke et al., 2019; Wolf et al., 2018) where a combination of a strong boundary predictor and graph partitioning methods has been shown to deliver accurate segmentation results. Briefly, the method consists of two major steps. In the first step, a convolutional neural network (CNN) is trained to predict cell boundaries. Then, a region adjacency graph is constructed from the pixels with edge weights computed from the boundary predictions. In the second step, a partitioning of the region adjacency graph is computed to produce the segmentation. The accuracy of this method is highly dependent on the boundary segmentation given by the CNN. Since no ground truth segmentation was initially available for our data, no dedicated CNN could be trained to accurately segment cell membranes. Instead, we used the following iterative procedure. In the first iteration a pre-trained CNN available in the PlantSeg package was used to generate the initial membrane probability maps. Specifically, we used a CNN trained on the confocal stacks of the Arabidopsis ovules dataset named “*confocal_unet_bce_dice_ds2*”. Having the cell boundary prediction, the initial segmentation was produced with PlantSeg. Then, the segmentation has been improved by visually choosing the most correctly segmented regions, manually correcting the remaining errors and using the results as training data for a dedicated neural network for the membrane prediction task. This process of choosing the best segmentation, proofreading and re-training the network was performed three times. A fourth optimisation was performed individually for each dataset, using 5 timepoints regularly spaced in the time series and manually curated to generate a dedicated neural network for the membrane prediction of a unique dataset, allowing for the segmentation of hundreds of timepoints with almost no curation required.

The resulting network was used as a basis for the graph partitioning in PlantSeg. For the final segmentation of embryos into cells we have chosen the MultiCut graph partitioning strategy (Andres et al., 2011; Bansal et al., 2004; Kappes et al., 2011), implemented (Pape et al., 2017) in PlantSeg. The best hyperparameters for the MultiCut algorithm have been found by a grid-search procedure which explored thousands of different configuration parameters simultaneously using EMBL’s computing cluster. Manual curation and editing were performed by one operator using a custom-made application (unpublished) and validated by a second operator.

### Preparation of the labelled data for analysis

After curation, the segmentation data were scaled without interpolation with Fiji (Schindelin et al., 2012) to generate isotropic voxel size of 0.416 × 0.416 × 0.416 *µ*m^3^. Polar bodies were automatically removed from the segmentation file based on volume size and blastomeres were automatically tracked using our weighted nearest neighbour algorithm. Finally, segmentation holes and gaps between cells were closed with a dilation of one voxel around each labelled cell. For each embryo, we checked sudden changes of cell volume and Jaccard index over time to identify and correct potential segmentation errors.

### Parametrisation with exponential splines

#### Short summary

Each cell has been parametrised independently using exponential splines. Assuming a continuous surface without holes, cells are modelled as deformations of a continuously-defined parametric spline sphere with *n* longitudes and *m* latitudes. In this study, we used *n=5* and *m=5* to maximise the accuracy of the geometrical approximation while avoiding over-fitting of artefactual information (e.g., voxel aliasing). Equations have been re-derived from previous work (Delgado-Gonzalo et al., 2013) to accommodate for mathematical errors. The Python package *splinefit* is available as part of the package *morphomap* (see section *Code availability*).

#### Setup

Data are assumed to be 3D image volumes (in .tif format) containing integer masks for each individual cells. Each cell is a single connected component which voxels are all labelled with the same integer value. Label values may be any unique integer, and background voxels are labelled as 0.

#### Fitting pipeline

Cells are assumed to have a sphere topology (no void, no hole). We therefore model them as deformations of a continuously-defined parametric spline sphere. The resulting closed surfaces are *C*^2^ smooth. We advocate for a continuously-defined parametric (implicit) representation as it allows for fast surface comparison across physical object size and voxel size. In this pipeline, each cell is processed independently. The embryo is thus defined as a collection of non-interacting deformed spheroids with no built-in notion of contact or spatial interactions.

#### Parametrisation, general form

The sphere **σ** of radius ***r*** ∈ ℝ, embedded in ℝ^3^, is defined in its parametric form for *s, t* ∈ [0,1] as

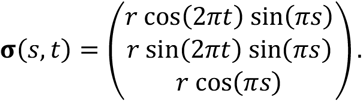

The two parameters, *t* and *s*, run along the latitudes and longitudes respectively, as depicted in Figure S2A. Note that latitudes are defined as full circles (closed), while longitudes are half-circle arcs (open). The parametric form of a 3D ellipsoid naturally follows for *s, t* ∈ [0,1] as

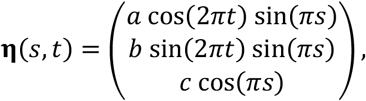

with *a, b, c* ∈ ℝ the three ellipse axes.

#### Parametrisation, spline model

The 3D ellipsoid **η** can be perfectly interpolated as a spline surface following

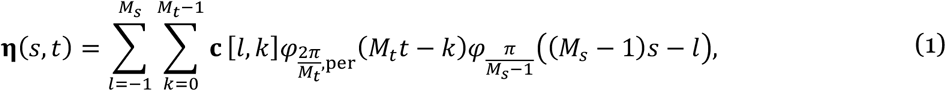

where **c**[*k, l*] are the parameters of the model, called control points. The integers *M*_*t*_ and *M*_***s***_ correspond to the number of parameters along each latitude and longitude, respectively. The function *φ*_*α*_: ℝ → ℝ is the exponential spline basis (Delgado-Gonzalo et al., 2012), given by

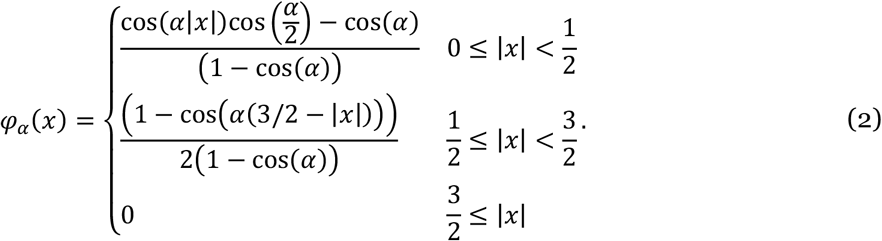

The basis *φ*_a=_ reproduces the space generated by {1, *x*, e^*jα*^, e^−*jα*^}. Because latitudes are closed circles, the basis associated to *t* is *M*_*t*_-periodized as

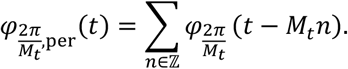

The expression (1) is here normalized such that the two continuous parameters run in [0,1].

For the ellipsoid to be properly closed, smoothness must be ensured at the poles. This translates to the two following conditions:

i. Interpolation, *i*.*e*., all longitudes meet at two well-defined points, the north pole **c**_N_ and the south pole **c**_S_:

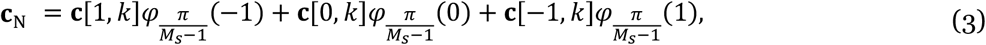

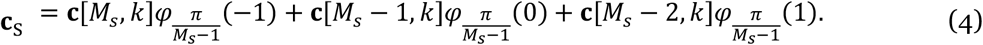
ii. Smoothness, *i*.*e*., each pole has a properly defined *tangent plane* characterized by two tangent plane vectors:

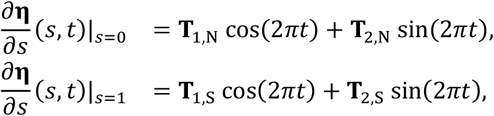

with **T**_1,N_ **T**_2,N_, **T**_1,S_, **T**_2,S_ ∈ ℝ^3^ the tangent plane vectors of the north and south pole, respectively. From (1), the above relations translate to

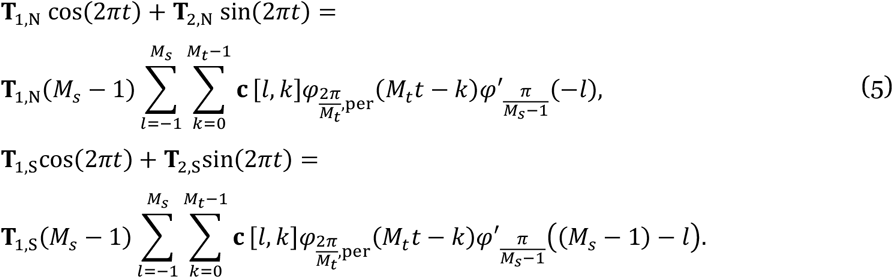

The initial set of control points parameterizing an ellipse of axes *a, b, c* ∈ ℝ and center **p**_0_ ∈ ℝ^3^, with north-south axis aligned along *z*, are given by

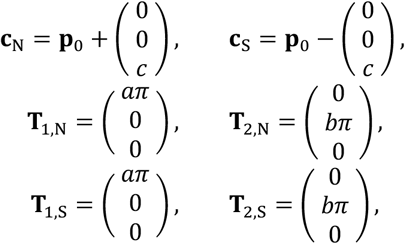

and

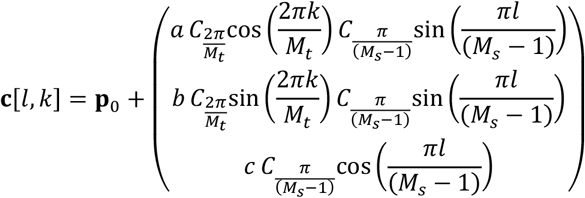

for *l* = 1, …, *M*_*s*_ − 2 and *k* = 0, …, *M*_*t*_ − 1, with

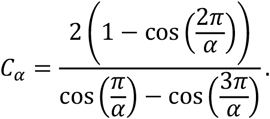

The surface (1) is thus fully determined by *M*_*t*_ (*M*_*s*_ − 2) + 6 parameters: **c**_N_, **c**_S_, **T**_1,N_, **T**_2,N_, **T**_1,S_, **T**_2,S_, and **c**[*l, k*] for *l* = 1, …, *M*_*s*_ − 2, *k* = 0, …, *M*_*t*_ − 1 (see also Figure S2B). Due to the support size of the exponential spline basis, extra control points are added at the extremities of each open longitude to ensure a correct behaviour at the boundaries. The extra control points **c**[*l, k*] for *l* = −1,0, *M*_*s*_ − 1, *M*_*s*_, *k* = 0, …, *M*_*t*_ − 1 are computed from the known parameters as

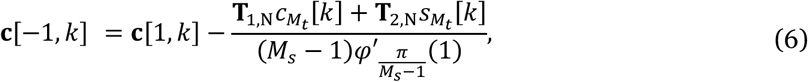

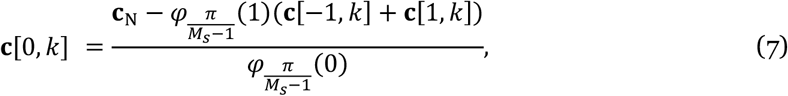

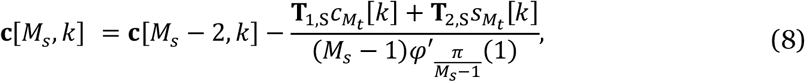

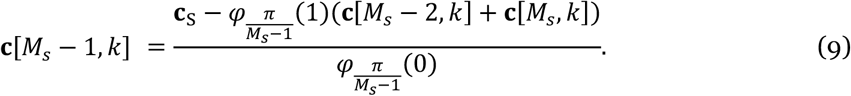

With

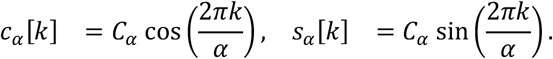

The relation (6) is obtained by identifying that

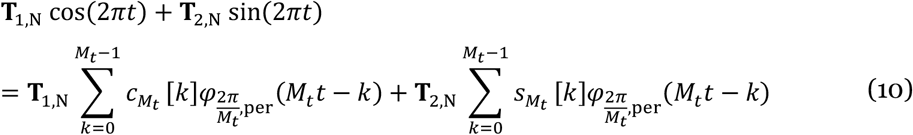

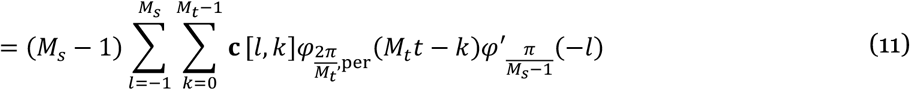

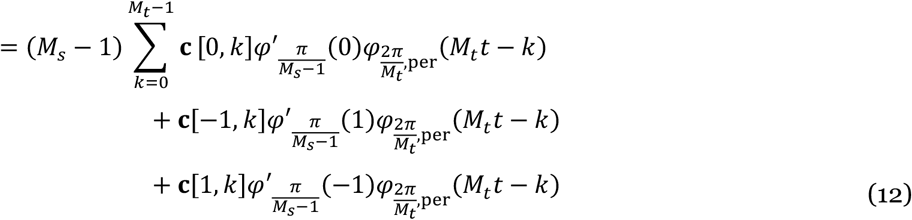

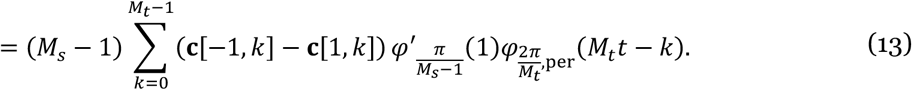

First, (10) is the direct expansion of the cosine and sine in the basis *φ*_*a*_ (2). Then, (11) is obtained directly from (5). Finally, (12) is obtained by noticing that, because *φ*_*a*_ is supported in 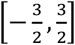, all terms in the sum vanish except for *j* = −1,0,1. The symmetry properties of the basis impose that *φ*′_*α*_(0) = 0 and *φ*′_*α*_ (−1) = −*φ*′_*α*_(1), yielding (13). One obtains (8) in a similar way. The relations (7) and (8) are obtained directly from (3) and (4), respectively. Note that **c**[0, *k*] ≠ **c**_N_ and **c**[*M*_*s*_, *k*] ≠ **c**_S_ due to the non-interpolatory behaviour of *φ*_*α*_.

For visualization purpose, the continuously-defined spline surface can be discretized as 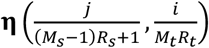 for *j* = 0, …, (*M*_*s*_ − 1)*R*_*s*_ and *i* = 0, …, *M*_*t*_*R*_*t*_ − 1, with *R*_*t*_, *R*_*s*_ ∈ ℕ* some user-defined sampling rates (*i*.*e*., amount of samples in each interval between successive control points) along *t* and *s*, respectively. In this study, we chose *M*_*s*_ = 5, *M*_*t*_ = 5, *R*_*s*_ = 5 and *R*_*t*_ = 5 to optimise smoothing, geometrical accuracy and computation time.

#### Axes identification

When parametrizing a cell, the identification of the north-south axis is an important design choice. We identified the following strategies:

i. Canonical ℝ^3^ basis: The north-south axis is aligned with *z* and control points are placed along rays 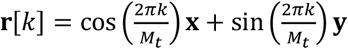 **y** in the *x, y* plane,
ii. Custom axis: We determine a rotation matrix **R** and a centre (e.g., obtained from the alignment of two embryos) onto which the model can be registered. The north-south axis is aligned with *z*′ = **R***z* and control points are placed along rays 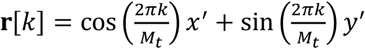 in the *x, y* plane, with *x*′ = **R***x* and *y*′ = **R***y*.

#### Data sampling, ray tracing

Considering the surface model (1) and well-defined parametrization axes, interpolation points are identified by ray tracing in order to reconstruct the spline surface from voxel data. Ray tracing is implemented relying on the 3D Digital Differential Analyzer (DDA) algorithm (Amanatides and Woo, 1987). A ray between two floating-point valued positions **p**_0_ and **p**_1_ is defined for *t* ≥ 0 as **p**_0_ + **v***t*, with **v** = **p**_1_ − **p**_0_. The algorithm is initialized by defining, for each image dimension *d*, the two quantities 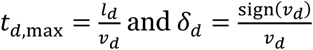. The DDA then proceeds as described by the following algorithm:

##### Result

Matrix **P** of grid intersection points

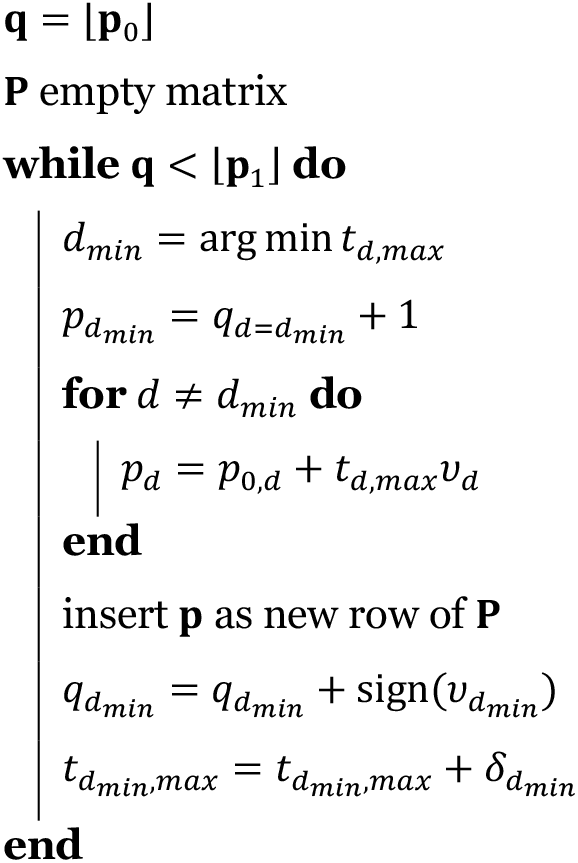

Each step of the algorithm requires *D* floating point comparisons (where *D* is the image dimension) and 2 floating point addition, making it reasonably fast.

In our case, data contain instance segmented volumes and ray tracing is thus carried out to identify sample points on the object surface. In this context, rays are traced in latitude planes, starting from the north-south axis and expanding until the image boundaries. Tracing stops at the first zero-labelled voxel.

#### Data sampling, interpolation

Given a set of data points 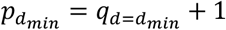 on the surface, the corresponding **c**[*l, k*] can be estimated by solving the linear system

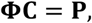

where

- **C** is a *M*_*t*_(*M*_*s*_ + 2) × 1 *control points matrix* with entries 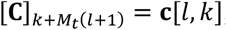,
- **Φ** is a (*M*_*t*_*R*_*t*_)((*M*_*s*_ − 1)*R*_*s*_ + 1) × *M*_*t*_(*M*_*s*_ + 2) *basis functions matrix* with entries 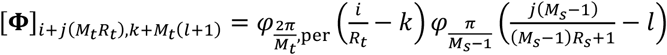
- **P** is a (*M*_*t*_*R*_*t*_)((*M*_*s*_ − 1)*R*_*s*_ + 1) × 1 *data point matrix* with entries 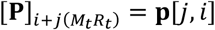..

The system is solved for **C** by finding the least-square best solution that minimizes the squared *l*_2_ norm 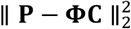. The poles are estimated from **c**[*l, k*], *l* = −1, …, *M*_*s*_,*k* = 0, …, *M*_*t*_ − 1, relying on (7) and (9) as

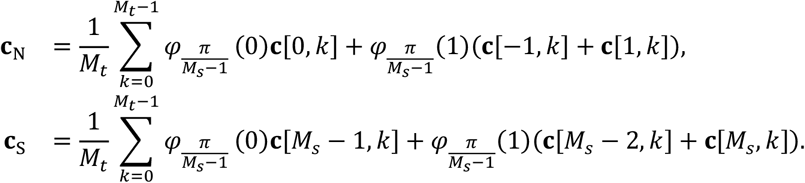

The tangent planes are retrieved from (6) and (8). For the north tangent plane,

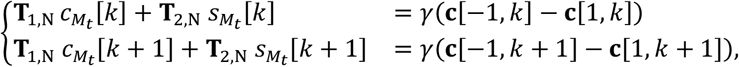

leading to

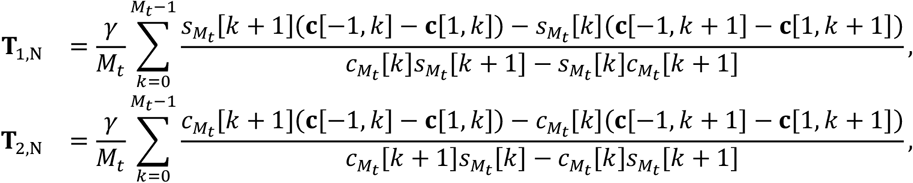

With

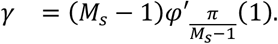

A similar development follows for the south tangent plane.

## Time normalisation at the 8-cell stage

Time progression through the 8-cell stage was normalised between 0 (first timepoint with 8 cells) and 1 (last timepoint with 8 cells).

## Pair-wise distance and 2D projection

To compute the geometrical distance between two sets of 3D image, we first generated the corresponding segmentation S_A_ and S_B_. Second, we used our nearest neighbour algorithm to determine the best transformation matrix (M) and the best list of cell assignment between S_A_ and S_B_. Then we computed the spline parameters for S_A_ with a North/South axis aligned with the Z-axis of the image, and the spline parameters for S_B_ with a North/South axis V such that the MV aligned with the Z-axis of the image. Finally, for each pair of assigned cells between S_A_ and S_B_, we computed the squared Euclidean distance (d) between spline parameters with the same latitude/longitude. The final geometrical distance was defined as the sum of all the distances d, divided by 10,000 for visual clarity. The projection was done using the implementation of the tSNE algorithm found in scikit-learn (Pedregosa et al., 2011). The morphomap pipeline was implemented in the Python package *morphomap* (see section *Code availability*).

## Measurement of the *a*-parameter

To determine the *α*-parameter, we used the segmentation data. We developed the Python package *interfaces* (see section *Code availability*) to determine the angle between two cells, and the corresponding value of *α*. First, we identified all the voxels on the surface of the embryo (V_S_, non-zero voxels with 6-connected neighbours of exactly 2 voxel types, itself and zero), all the voxels on the external line interface between two cells (V_L_, non-zero voxels with 6-connected neighbours of exactly 3 different voxel types, itself, zero and a third arbitrary value) and all the voxels belonging to the external interface between three cells or more (V_P_, non-zero voxels with 6-connected neighbours of at least 4 different voxels types, itself, zero and other arbitrary values). We defined two parameters: the exclusion radius R_E_ and the inclusion radius R_I_.

For each points P of V_L_, between cell A and B, that is further than R_E_ away from any points in V_P_, we listed all the points P_A_ and P_B_ of V_S_ that are within a R_I_ distance from P and belongs to cell A and B respectively. We fitted two plans such that P belongs to them and the Euclidean distance to P_A_ and P_B_ is minimised. Finally, we computed the angle between the two plans and determined the *α*-parameter using the Young-Dupré equation as previously described (Maître et al., 2015). R_E_ (30 voxels) and R_I_ (15 voxels) values were optimised to accurately described theoretical data with known *α*-parameter.

## Estimation of contacts and surface areas

To estimate the surface area, we used our Python package *interfaces* (see section *Code availability*) and we triangulated the voxels belonging to the surface of interest using a marching cube algorithm (Lewiner et al., 2003) and the surface area measurement implemented in scikit-image (van der Walt et al., 2014). Using the segmentation data, the interface between two cells was defined as the list of non-zero voxels with 6-connected neighbours of exactly 2 voxel types. Note that the interface between a cell and the background is included (one voxel type being zero). The triangulation was performed on the whole cells, but only the triangles less than 2 voxels away from the interface were used in the estimation of the surface area. The interface surface areas were linearly corrected to ensure that the sum of all the interface surface areas equals the estimated surface of the whole cell.

## Identification of the closest rigid packing

### Preliminaries

i. We refer to the 13 rigid packings possible with 8 hard spheres described in Arkus et al., 2009 as the set of the 13 ideal packings,
ii. Data of cell contacts is provided as a matrix where the two indices of each position represent the labelling of the two cells in potential contact,
iii. The entries of the matrix display the proportion of the surface area of the first cell involved in the contact with the second cell.

### Rigidity condition

Prior to any computation, we perform a first check whether the topological structure under study is rigid or floppy. In rigid structures, no independent movements of the elements are allowed unless we perform some work over the system (Jacobs, 1998) (see also Figure S3A). For three dimensions, the condition of rigidity for a system with *n* elements and *k* constraints (in our case, links) states that:

a. *k* ≥ 3*n* − 6 which, in the case of *n* = 8 implies that *k* ≥ 18 that is, that the network of cell cell-contacts has at least 18 links,
b. No node of the network has less than three links. That means that no cell of the packing is in contact to less than 3 other cells.

### Distance between two arbitrary packings

To evaluate the distance between two packings we must consider the adjacency matrix of the network describing the topological structure of the packing. Given a packing made of *V* cells (*ν*_1_, …, *ν*_8_), the values of the adjacency matrix *A* of the packing are defined such that *A*_*ij*-_ = 1 if cell *ν*_*i*_ and *ν*_*j*_ are in contact and *A*_*ij*_ = 0 otherwise. Given that there are 8! = 40320 different ways to label the cells leading to the same topological structure, the direct comparison of adjacency matrices is not affordable. Furthermore, one must consider that soft spheres may define contacts in places where hard spheres cannot, making the space of possible equivalent configurations even bigger. Therefore, it is much more suitable to work with the ordered spectrum of eigenvalues, which is invariant under permutation of the adjacency matrices. Since, by construction, the adjacency matrix is symmetric and made of non-negative real entries, the eigenvalue spectrum is all made of real numbers. Thanks to this last property, one can easily compute the distance between two packings: The vector representing the eigenvalue spectrum of the adjacency matrix *A* is defined as *u*_*A*_ = (*λ*_1_, *…, λ*_8_) where *λ*_1_ > *λ*_2_ > … > *λ*_8_. Consider that these two packings can be represented by adjacency matrices *A* and *A*′, respectively. The distance between these two packings is obtained by considering the Euclidean distance between the vectors representing the (ordered) eigenvalue spectrum of each:

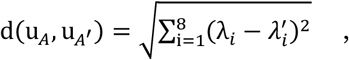

where

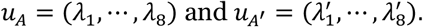

### Finding the closest ideal packing to the real embryo packing

To identify the closest ideal packing to the topological structure of the embryo under study, we proceed as follows:

i. Check if the number of contacts of the embryo is 18 –recall that we already filtered the non-rigid packings,
ii. If the number of contacts is larger than 18, remove those with the lowest contact area until we reach the exact number of 18 contacts,
iii. Binarize the matrix of contacts such that all contacts are set to 1,
iv. Compute the distance to all 13 ideal rigid packings as described above.

Pick the ideal packing displaying the closest distance

## Enumeration of topological transitions

A topological transition was defined as two consecutive timepoints with different identified closest minimally rigid packing. Temporary transitions lasting one timepoint were ignored from the analysis unless the identified topologies right before and right after were different. For example, the sequence AABABB (3 transitions, with A and B two arbitrary topologies) was transformed to AAAABB (one transition), while the sequence AABCC (2 transitions, with C a third arbitrary topology) was not modified.

## Measurement of the packing parameter

We first determine the centroid *c*_*i*_ of the outer cells *i*. We then used the average coordinates of outer cells’ centroid to obtain the centroid of the embryo *e*. For every outer cell, we computed the Euclidean distance *d*_*i*_ from *c*_*i*_ to *e*. The packing parameter was defined as the invert of the standard deviation of *{d*_*1*_, *d*_*2*_, *…, d*_*i*_*}* (see Figure S6a for a graphical representation of the packing parameter).

## Inner/outer cell classification

Cells at the 8-cell stage were all considered outer. For the 16-cell stage, we computed the contact-free area (i.e., the surface area of the cell that is not in contact with another cell) and binned the cells in 10 groups from 0% (completely inside) to approximately 60% (most of the cell surface exposed to the outside environment). The resulting count histogram exhibited a bi-modal distribution. We fitted the histogram values with the sum of two weighted Gaussian distributions. The cut-off between inner and outer cells was defined as the value of contact-free surface area at which the two gaussians intersect.

## Count of ectopic Sox2^+^ and Cdx2^+^ cells

For the early blastocyst stage, fate-based cell type was determined as Sox2^+^ (Cdx2^+^ respectively) if the immunostaining signal intensity for Sox2 (Cdx2) was higher than the one for Cdx2 (Sox2). Sox2^+^ and Cdx2^+^ cells were labelled as *inner* and *outer* respectively, based on fate markers. Additionally, we computed the *α*-shape of the cell centres (Edelsbrunner and Mücke, 1994) using the R package *alphashape3d* (Lafarge and Pateiro-Lopez, 2020) and labelled the cells as *outer* and *inner* if they belongs to the surface determined by the *α*-shape or not respectively. Ectopic cells are, by definition, cells with label discrepancies between fate markers and their positions in the embryo. Embryos with no identified *inner* or *outer* cells based on marker signal intensity and/or cell coordinates were excluded from the analysis.

## Dataset usage

Use of live imaging datasets in figures is described in Table S2.

## Code availability

Code used to produce the figure panels and the analyses in this study is available on the following public repository: https://gitlab.com/fabreges/fabreges-2023/

## Statistical Analysis

All statistical analyses were performed using RStudio 2022.12.0+353 (Posit team, 2022) with R 4.2.2 (R Core Team, 2022), with the base library and the following libraries: *alphashape3d* (Lafarge and Pateiro-Lopez, 2020), *ggplot2* (Wickham, 2016), *ggrepel* (Slowikowski et al., 2022) and *namespace* (Chang et al., 2012). No statistical analysis was used to predetermine sample size. No sample was excluded. No randomization method was used. The investigators were not blinded during experiments. Sample sizes, statistical tests and p-values are indicated in the text, figures and figure legends. n-values indicate number of embryos analysed for different experimental conditions unless mentioned otherwise. Error bars indicate mean ± s.d. unless mentioned otherwise.

