## Supplemental figures and legends for "Temporal variability and cell mechanics control robustness in mammalian embryogenesis"

### Figure S1

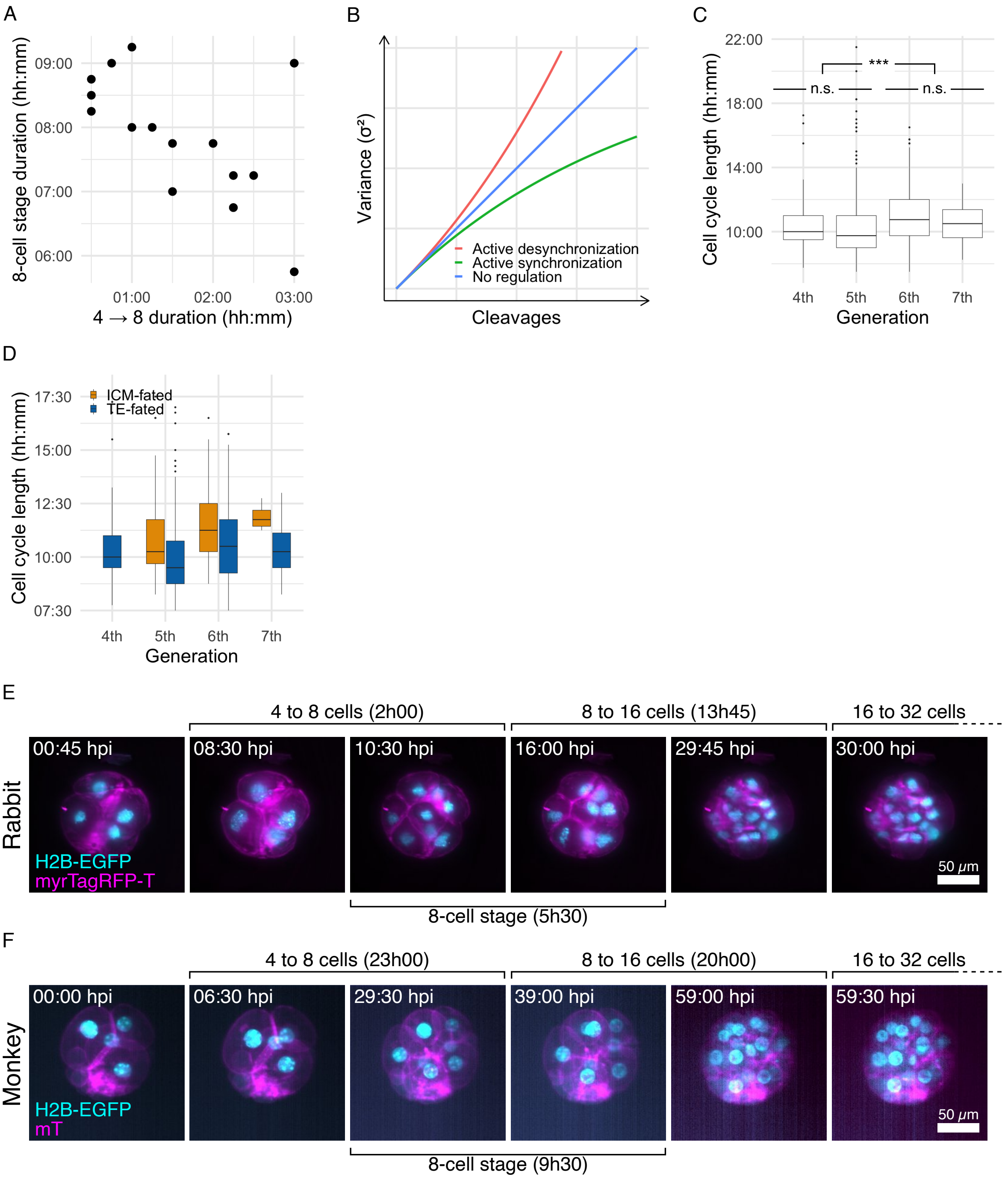

A
8-cell stage duration (hh:mm)
09:00
08:00
07:00
06:00
01:00
02:00
03:00
4 → 8 duration (hh:mm)
B
Variance (σ²)
Cleavages
Active desynchronization
Active synchronization
No regulation
C
Cell cycle length (hh:mm)
22:00
18:00
14:00
10:00
***
n.s.
n.s.
4th
5th
6th
7th
Generation
D
Cell cycle length (hh:mm)
17:30
15:00
12:30
10:00
07:30
ICM-fated
TE-fated
4th
5th
6th
7th
Generation
E
4 to 8 cells (2h00)
8 to 16 cells (13h45)
16 to 32 cells
Rabbit
00:45 hpi
08:30 hpi
10:30 hpi
16:00 hpi
29:45 hpi
30:00 hpi
H2B-EGFP
myrTagRFP-T
50 μm
8-cell stage (5h30)
F
4 to 8 cells (23h00)
8 to 16 cells (20h00)
16 to 32 cells
Monkey
00:00 hpi
06:30 hpi
29:30 hpi
39:00 hpi
59:00 hpi
59:30 hpi
H2B-EGFP
mT
50 μm
8-cell stage (9h30)

**Figure S1. Embryo variability in cleavage timing increases at a constant and species-specific rate .**

(**A**) Duration of the 8-cell stage inter-mitotic period as a function of the duration of the 3rd cleavage from 4 to 8 cells. (**B**) Cell cycle length by cell generation. n.s., non-significant. ***, p-value < 0.001. (**C**) Cell cycle length by cell generation and prospective cell type. ICM-fated (orange) and TE-fated (blue) cells have been determined by retro-tracking of $Sox2^+$ and $Cdx2^+$ cells identified from immunostaining of the embryos fixed less than 15 minutes after the end of the imaging. (**D**) Schematic representation of three possible desynchronization dynamics. Blue, no regulation. Red (top curve), active mechanism increasing desynchronization. Green (bottom curve), active mechanism decreasing desynchronization. (**E,F**) Maximum projection of a representative live imaging of monkey (E) and rabbit (F) embryo expressing mT (E) or myrTagRFP-T (F) (magenta); H2B-EGFP (cyan) developing from the 4- to the 16-cell stage. Time post imaging (hh:mm); Scale bar, 25 $\mu$m. See also Video S2 and S3 respectively.

### Figure S2

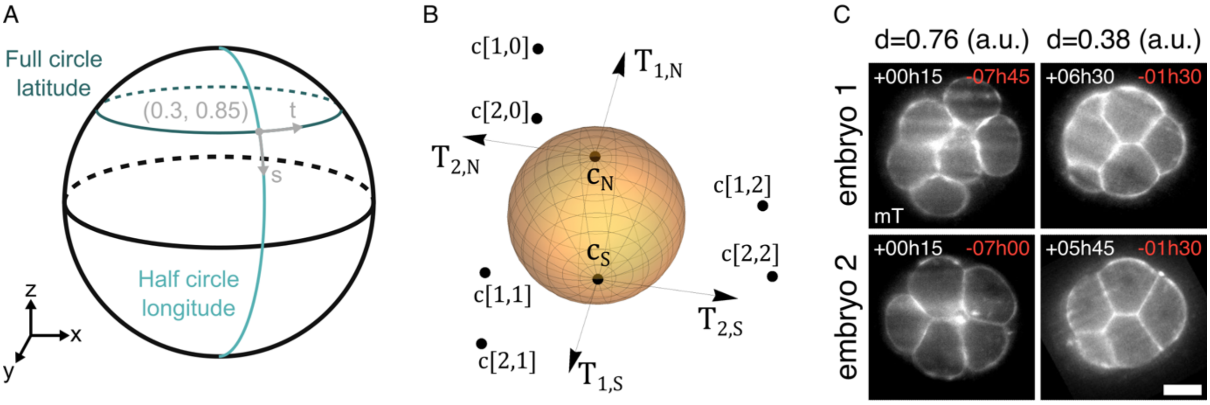
A
Full circle
latitude
(0.3, 0.85)
t
s
Half circle
longitude
z
x
y
B
c[1,0]
c[2,0]
$T_{1,N}$
$T_{2,N}$
$c_N$
$c_S$
c[1,2]
c[2,2]
c[1,1]
c[2,1]
$T_{2,S}$
$T_{1,S}$
C
d=0.76 (a.u.)
d=0.38 (a.u.)
embryo 1
+00h15
-07h45
+06h30
-01h30
mT
embryo 2
+00h15
-07h00
+05h45
-01h30

D

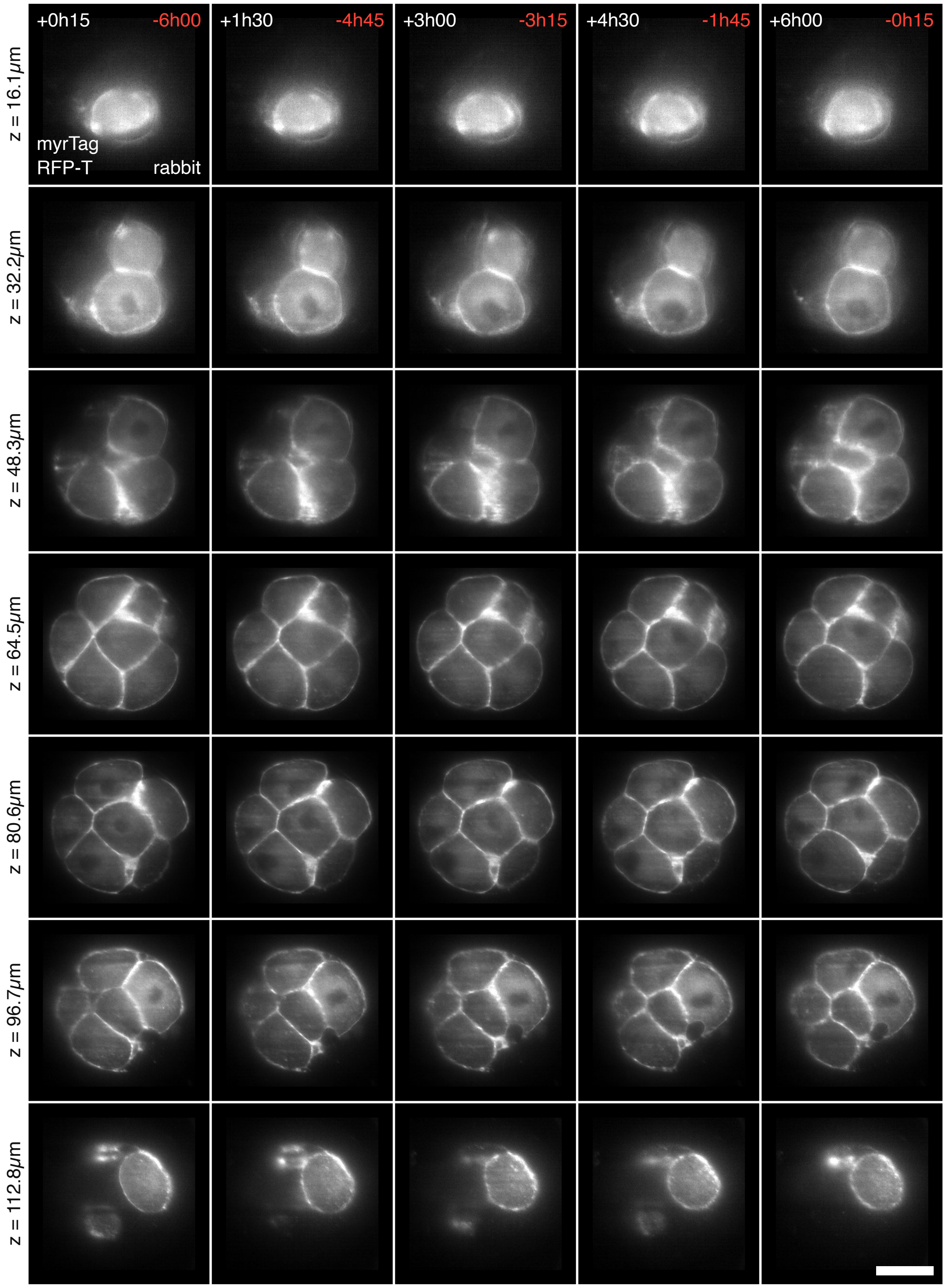
+0h15
-6h00
+1h30
-4h45
+3h00
-3h15
+4h30
-1h45
+6h00
-0h15
z = 16.1μm
z = 32.2μm
z = 48.3μm
z = 64.5μm
z = 80.6μm
z = 96.7μm
z = 112.8μm
myrTag
RFP-T
rabbit

E

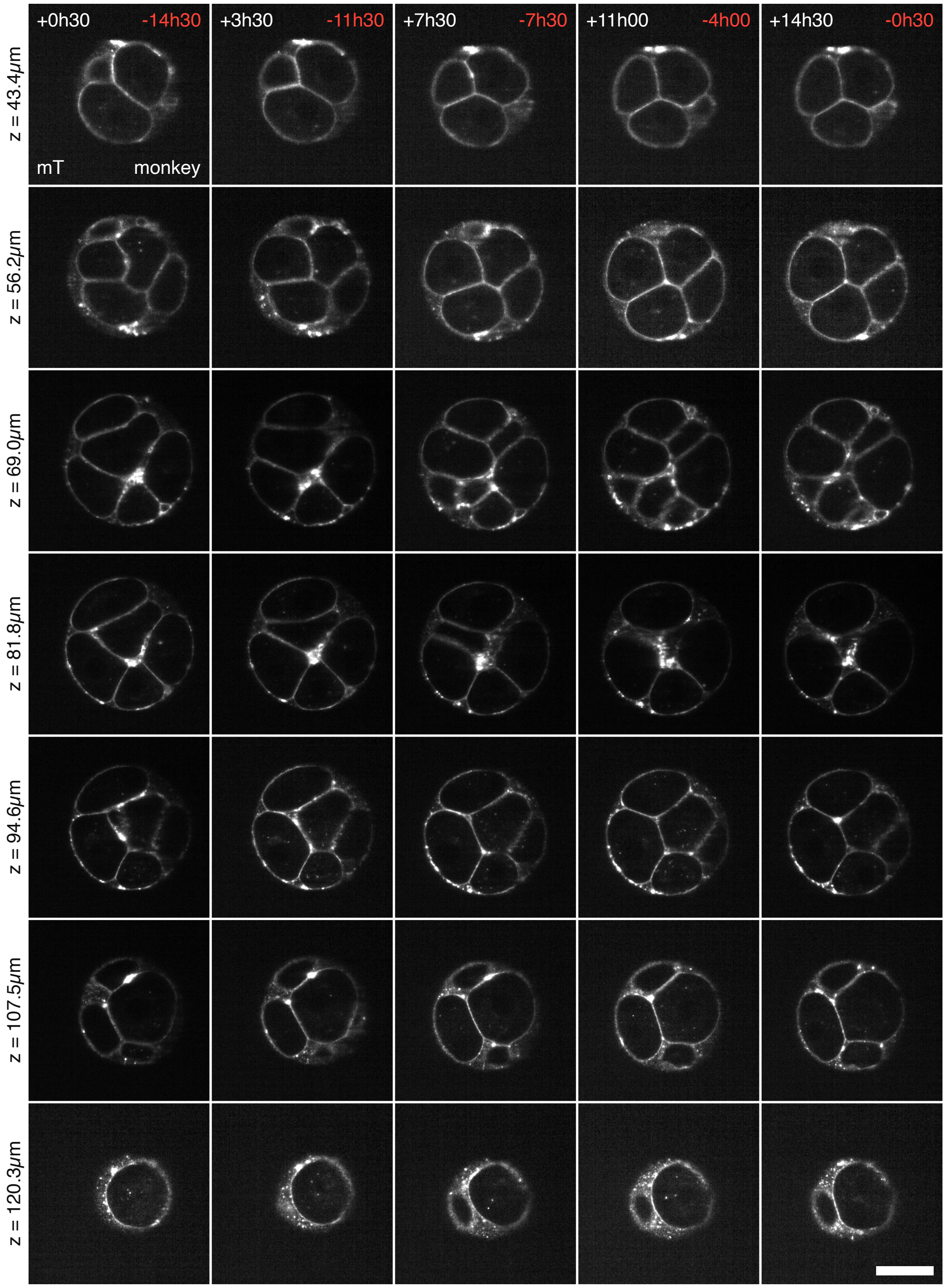

+0h30
-14h30
+3h30
-11h30
+7h30
-7h30
+11h00
-4h00
+14h30
-0h30
mT
monkey
z = 43.4μm
z = 56.2μm
z = 69.0μm
z = 81.8μm
z = 94.6μm
z = 107.5μm
z = 120.3μm

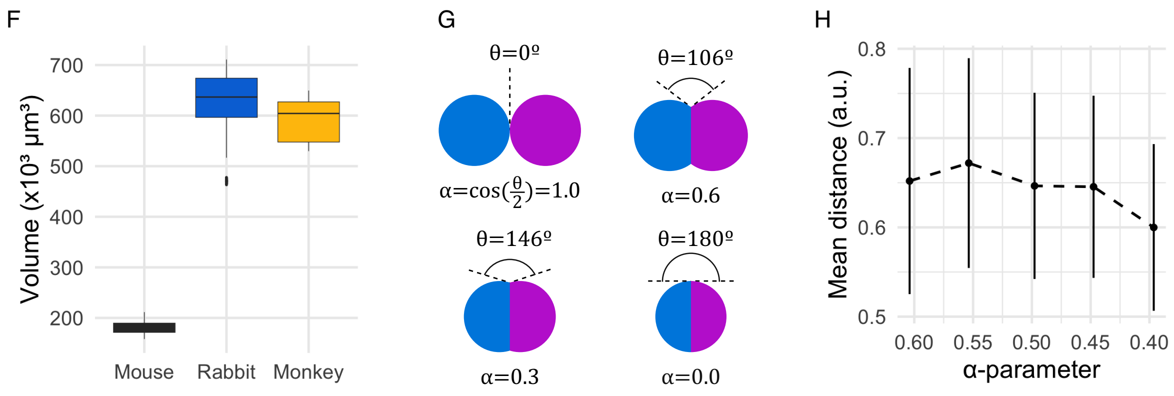
F
Volume (x10³ µm³)
700
600
500
400
300
200
Mouse
Rabbit
Monkey
G
θ=0º
α=cos(θ/2)=1.0
θ=106º
α=0.6
θ=146º
α=0.3
θ=180º
α=0.0
H
Mean distance (a.u.)
0.8
0.7
0.6
0.5
0.60
0.55
0.50
0.45
0.40
α-parameter

**Figure S2. Embryo spatial variability reduces during the 8-cell stage.**

(**A**) Parametric representation of the sphere of radius $r$ in $\mathbb{R}^3$ as used by the exponential spline to the surface of the cells (see also Methods). The latitudes correspond to full circles in the XY plane, parametrised as $(cos(2\pi t),\ sin(2\pi t), 1)$. The longitudes correspond to half-circle arcs along the Z axis, parametrised as $(sin(\pi s),\ sin(\pi s),\ cos(\pi s))$. A point located at ($s = 0.3$, $t = 0.85$) on the sphere is depicted as example. (**B**) A parametric spline ellipsoid is entirely determined by $Mt(Ms\ - 2) + 6$ parameters: a north pole $\boldsymbol{c}_N$ and its tangent plane defined by the two vectors $\boldsymbol{T}_{1,N}$, $\boldsymbol{T}_{2,N}$, a south pole $\boldsymbol{c}_S$ and its tangent plane defined by the two vectors $\boldsymbol{T}_{1,S}$, $\boldsymbol{T}_{2,S}$, and a set of control points $\boldsymbol{c}[l, k]$, with $l = 1, \dots, M_s$ and $k = 0, \dots, M_t - 1$ (see also Methods). The spline sphere with $M_s = 4$ and $M_t = 3$ is depicted as example. (**C**) Representative cross-section of two mouse embryos at the beginning (left column) and the end (right column) of the 8-cell stage, illustrating the geometrical convergence from distance 0.76 to 0.38 respectively (arbitrary unit). Time in hours after the beginning (white, top-left corner) and before the end (red, top-right corner) of the 8-cell stage. Scale bar, 25 $\mu$m. (**B,C**) Cross-sections of a representative live imaging of a rabbit (B) and a monkey (C) embryo during the 8-cell stage. Left to right, time in hours after the beginning (white, top-left corner) and before the end (red, top-right corner) of the 8-cell stage. Top to bottom, depth in $\mu$m. Scale bar, 50 $\mu$m. (**D**) Box plot representation of the total embryo volume in mouse (black, $n = 29$ embryos), rabbit (blue, $n = 10$ embryos) and monkey (yellow, $n = 4$ embryos). (**E**) Schematic representation of the link between compaction, cell-cell angle and the $\alpha$-parameter between two theoretical cells (magenta and cyan) with $\alpha = 1.0$ (no compaction), $\alpha = 0.6$, $\alpha = 0.3$ and $\alpha = 0.0$ (total compaction). (**F**) Mean ± s.d. of the pair-wise geometrical distances as a function of the $\alpha$-parameter in 29 mouse embryos.

### Figure S3

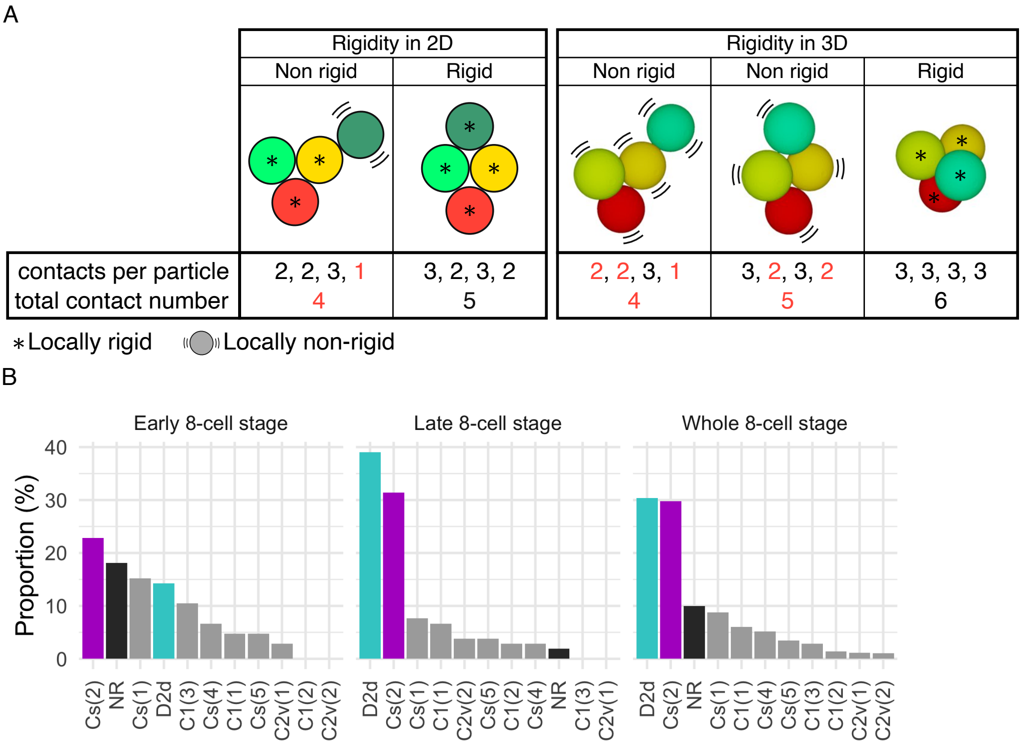

A
Rigidity in 2D
Non rigid
Rigid
Rigidity in 3D
Non rigid
Non rigid
Rigid
contacts per particle
total contact number
2, 2, 3, 1
4
3, 2, 3, 2
5
2, 2, 3, 1
4
3, 2, 3, 2
5
3, 3, 3, 3
6
* Locally rigid
Locally non-rigid
B
Early 8-cell stage
Late 8-cell stage
Whole 8-cell stage
Proportion (%)
40
30
20
10
0
Cs(2) NR Cs(1) D2d C1(3) Cs(4) C1(1) Cs(5) C2v(1) C1(2) C2v(2)
D2d Cs(2) Cs(1) C1(1) C2v(2) Cs(5) C1(2) Cs(4) NR C1(3) C2v(1)
D2d Cs(2) NR Cs(1) C1(1) Cs(4) Cs(5) C1(3) C1(2) C2v(1) C2v(2)

**Figure S3. Embryo topological variability reduces during the 8-cell stage**
(**A**) Illustration of the concept of structural rigidity in 2D and 3D for a cluster of 4 hard spheres. By definition, a cluster is rigid if and only if there are at least 2 (2D) and 3 (3D) contacts per sphere, and the total number of contacts is at least 5 (2D) or 6 (3D). No single sphere in a rigid packing can be moved relative to the other without an associated energy cost, while in non-rigid packings, cells can move without changing cell-cell contacts, thus the total energy (e.g., in 2D, the locally non-rigid circle in dark green can move freely around the adjacent yellow circle, while the locally rigid circles can only move by changing their connectivity to adjacent circles). For 8 spheres in 3D, a packing must have at least 18 contacts to be rigid. Numbers in red do not meet the conditions for the cluster to be rigid.
(**B**) Distribution of the proportion of $C_s(2)$ (magenta), $D_{2d}$ (cyan), non-rigid packings (NR, black) or other minimally rigid packings (grey) identified in mouse embryos during the first 10% (left), the last 10% (middle) and during the entire period (right) of the 8-cell stage ($n = 29$ embryos).

### Figure S4

A

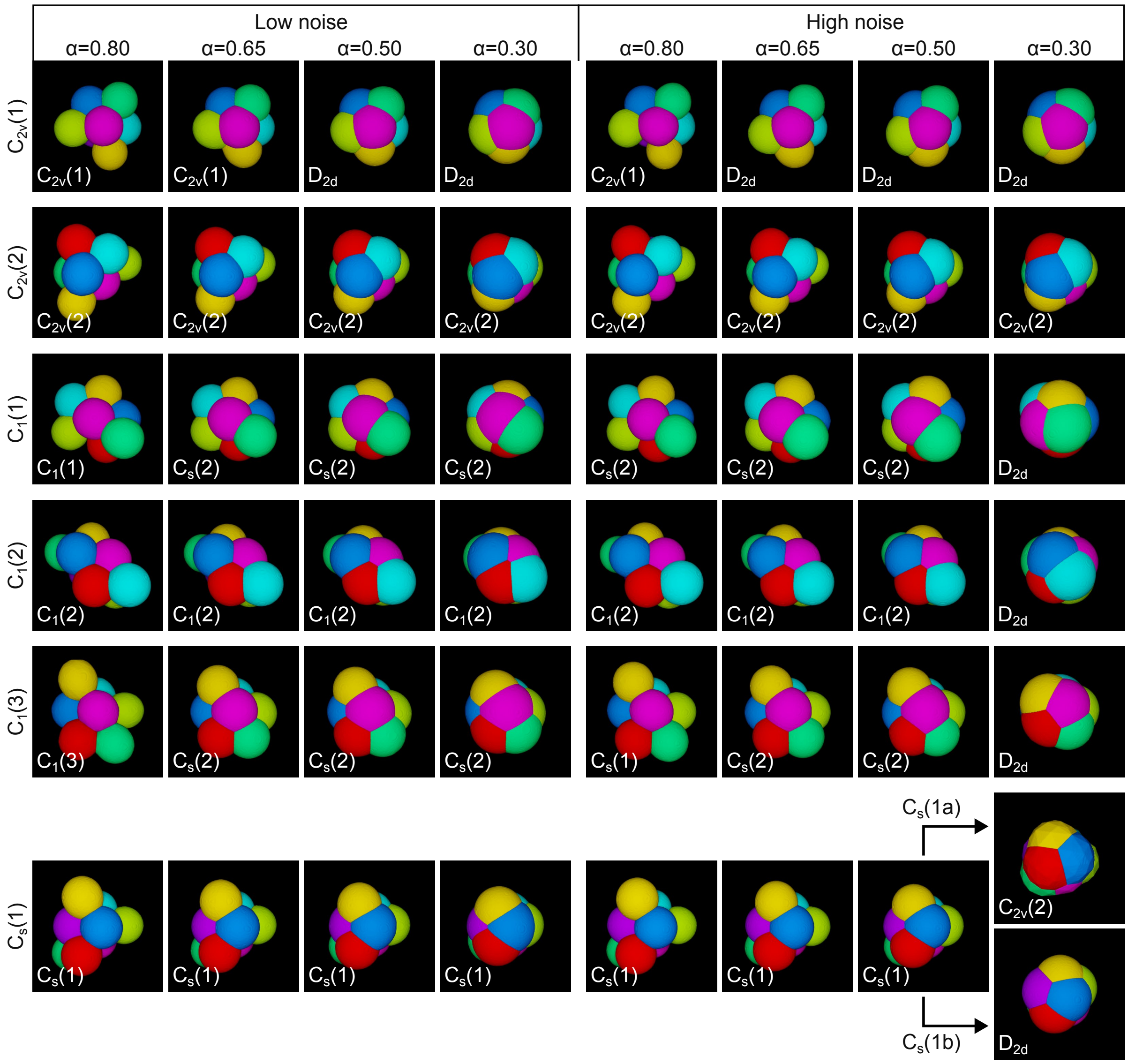
Low noise
High noise
α=0.80
α=0.65
α=0.50
α=0.30
α=0.80
α=0.65
α=0.50
α=0.30
C2v(1)
C2v(2)
C1(1)
C1(2)
C1(3)
Cs(1)
C2v(1) C2v(1) D2d D2d C2v(1) D2d D2d D2d
C2v(2) C2v(2) C2v(2) C2v(2) C2v(2) C2v(2) C2v(2) C2v(2)
C1(1) Cs(2) Cs(2) Cs(2) Cs(2) Cs(2) Cs(2) D2d
C1(2) C1(2) C1(2) C1(2) C1(2) C1(2) C1(2) D2d
C1(3) Cs(2) Cs(2) Cs(2) Cs(1) Cs(2) Cs(2) D2d
Cs(1) Cs(1) Cs(1) Cs(1) Cs(1) Cs(1) Cs(1)
Cs(1a)
C2v(2)
Cs(1b)
D2d

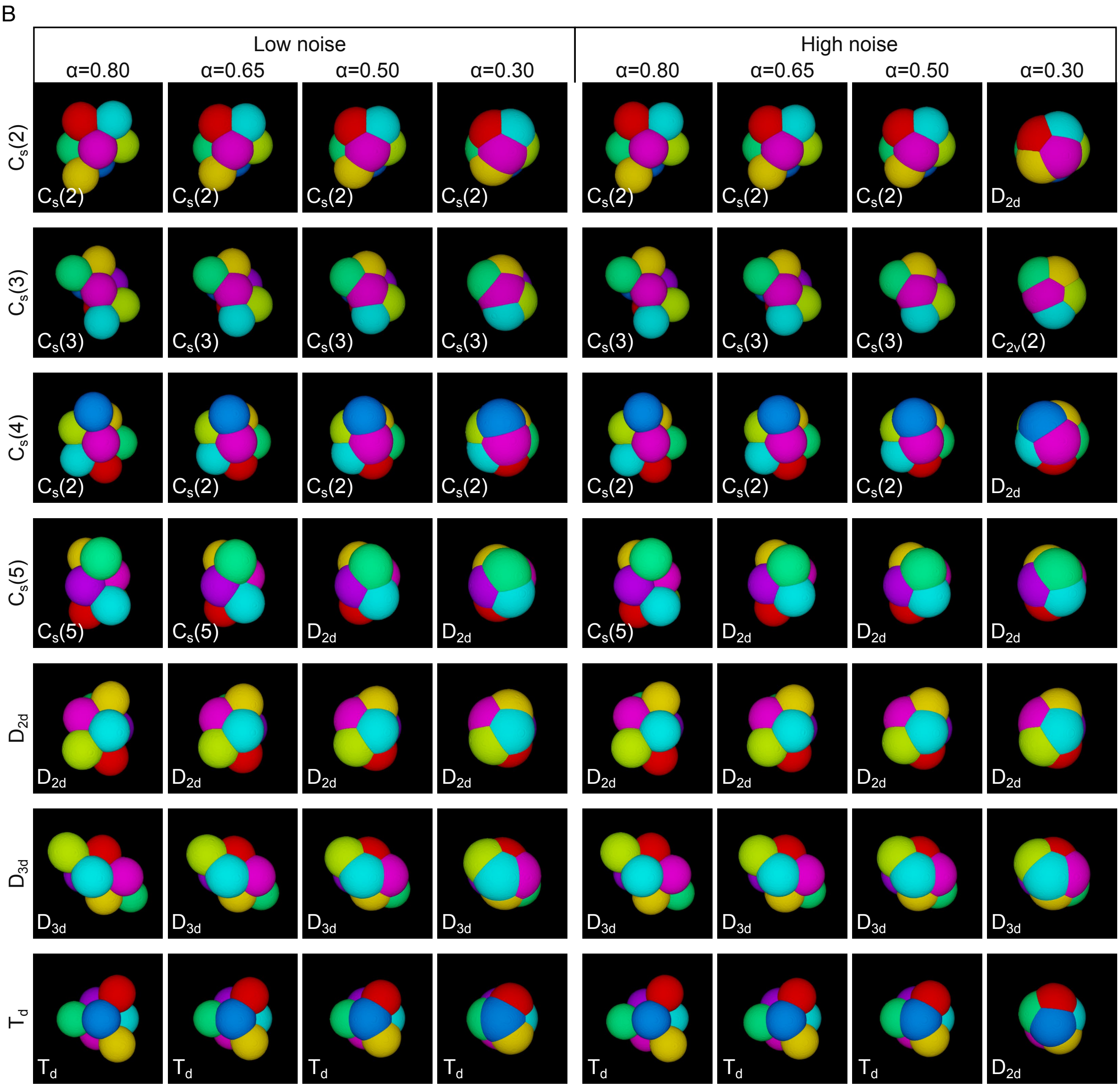

B
Low noise
High noise
α=0.80
α=0.65
α=0.50
α=0.30
α=0.80
α=0.65
α=0.50
α=0.30
Cs(2)
Cs(3)
Cs(4)
Cs(5)
D2d
D3d
Td
Cs(2) Cs(2) Cs(2) Cs(2) Cs(2) Cs(2) Cs(2) D2d
Cs(3) Cs(3) Cs(3) Cs(3) Cs(3) Cs(3) Cs(3) C2v(2)
Cs(2) Cs(2) Cs(2) Cs(2) Cs(2) Cs(2) Cs(2) D2d
Cs(5) Cs(5) D2d D2d Cs(5) D2d D2d D2d
D2d D2d D2d D2d D2d D2d D2d D2d
D3d D3d D3d D3d D3d D3d D3d D3d
Td Td Td Td Td Td Td D2d

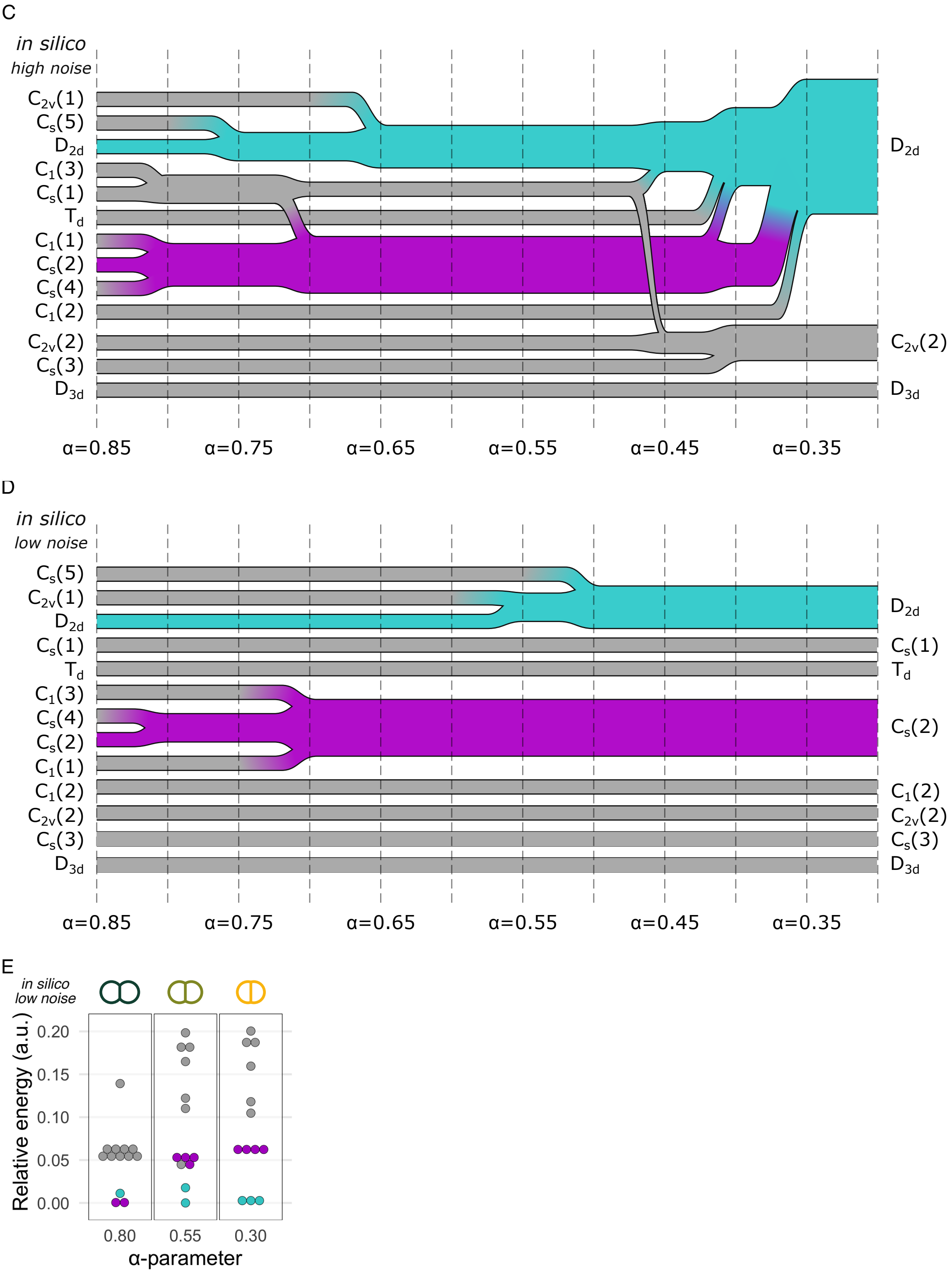
C
in silico
high noise
C2v(1)
Cs(5)
D2d
C1(3)
Cs(1)
Td
C1(1)
Cs(2)
Cs(4)
C1(2)
C2v(2)
Cs(3)
D3d
D2d
C2v(2)
D3d
α=0.85
α=0.75
α=0.65
α=0.55
α=0.45
α=0.35
D
in silico
low noise
Cs(5)
C2v(1)
D2d
Cs(1)
Td
C1(3)
Cs(4)
Cs(2)
C1(1)
C1(2)
C2v(2)
Cs(3)
D3d
D2d
Cs(1)
Td
Cs(2)
C1(2)
C2v(2)
Cs(3)
D3d
α=0.85
α=0.75
α=0.65
α=0.55
α=0.45
α=0.35
E
in silico
low noise
Relative energy (a.u.)
0.20
0.15
0.10
0.05
0.00
0.80
0.55
0.30
α-parameter

**Figure S4. Surface energy minimisation with compaction is sufficient to recapitulate geometrical and topological convergence.**
(**A,B**) Simulation of the compaction of the 13 rigid packings (rows) from $\alpha = 0.8$ to $\alpha = 0.3$ with low (left four columns) and higher (right four columns) level of noise. The initial condition used for the simulation is indicated at the beginning of each row. The closest minimally rigid packing is indicated in the lower left corner of each picture. Note that because the identification of the closest topology depends on the strength of the contacts between cells, interfaces of low surface area may have been discarded. $C_s(1)$ compaction with higher level of noise (A, bottom row) showed two possible topological transitions, towards $C_{2v}(2)$ and $D_{2d}$. (**C,D**) Topological transition flow of the 13 minimally rigid packings through progressive reduction of the $\alpha$-parameter *in silico* with low (B) and higher (C) level of noise, from and to $C_s(2)$ (magenta), $D_{2d}$ (cyan) and other rigid packings (grey). Stream thickness represents the proportion of topologies within each value of $\alpha$. (**E**) Relative energy of the packings obtained computationally after compaction of the 13 rigid packings to $\alpha = 0.8$, $0.55$ and $0.3$ with low noise. Colour code corresponds to the topological proximity to $C_s(2)$ (magenta), $D_{2d}$ (cyan) and other rigid packings (grey), and may differ from the initial packing used in the simulation due to topological transitions. See also Figure 4D.

### Figure S5

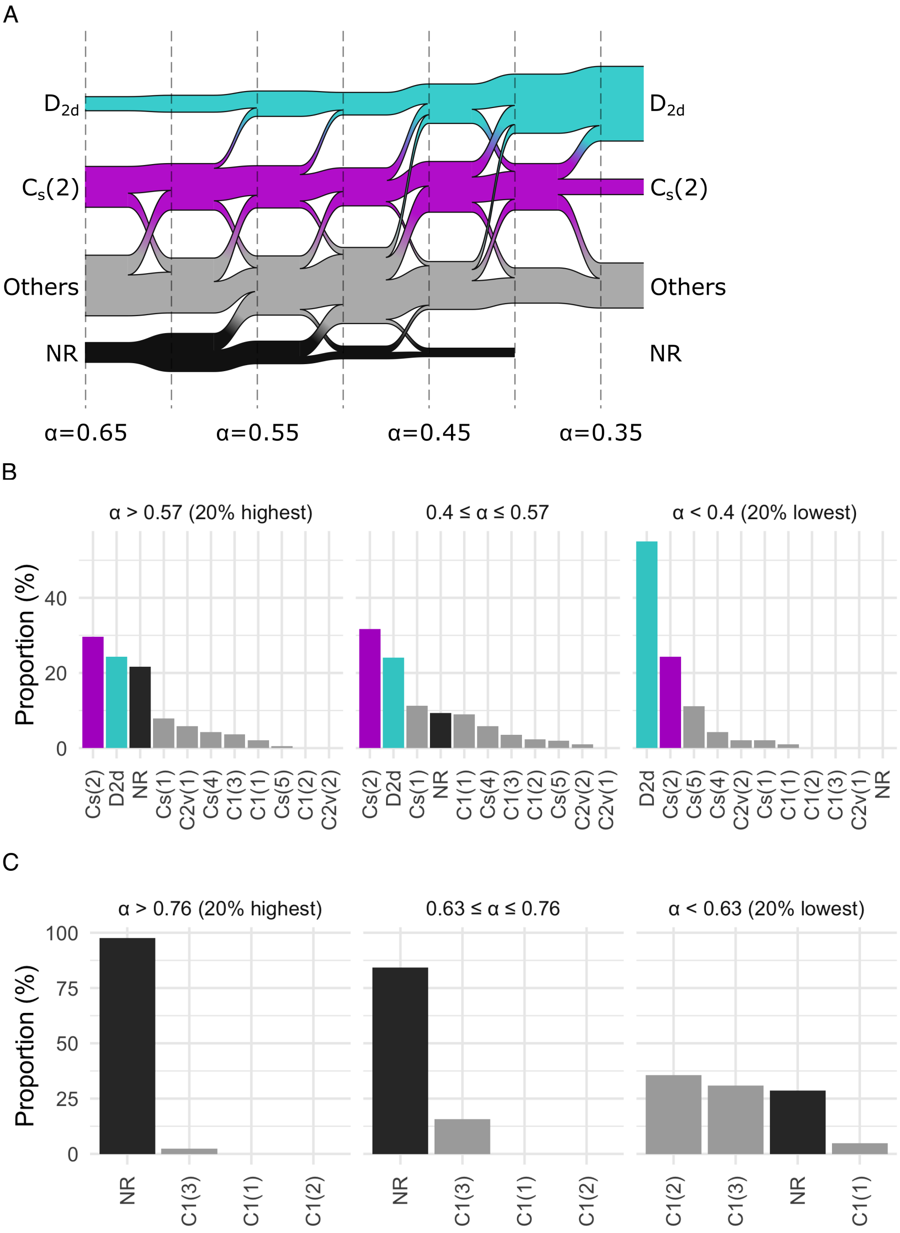
A
D2d
Cs(2)
Others
NR
α=0.65
α=0.55
α=0.45
α=0.35
B
α > 0.57 (20% highest)
0.4 ≤ α ≤ 0.57
α < 0.4 (20% lowest)
Proportion (%)
0
20
40
Cs(2) D2d NR Cs(1) C2v(1) Cs(4) C1(3) C1(1) Cs(5) C1(2) C2v(2)
Cs(2) D2d Cs(1) NR C1(1) Cs(4) C1(3) C1(2) Cs(5) C2v(2) C2v(1)
D2d Cs(2) Cs(5) Cs(4) C2v(2) Cs(1) C1(1) C1(2) C1(3) C2v(1) NR
C
α > 0.76 (20% highest)
0.63 ≤ α ≤ 0.76
α < 0.63 (20% lowest)
Proportion (%)
0
25
50
75
100
NR C1(3) C1(1) C1(2)
NR C1(3) C1(1) C1(2)
C1(2) C1(3) NR C1(1)

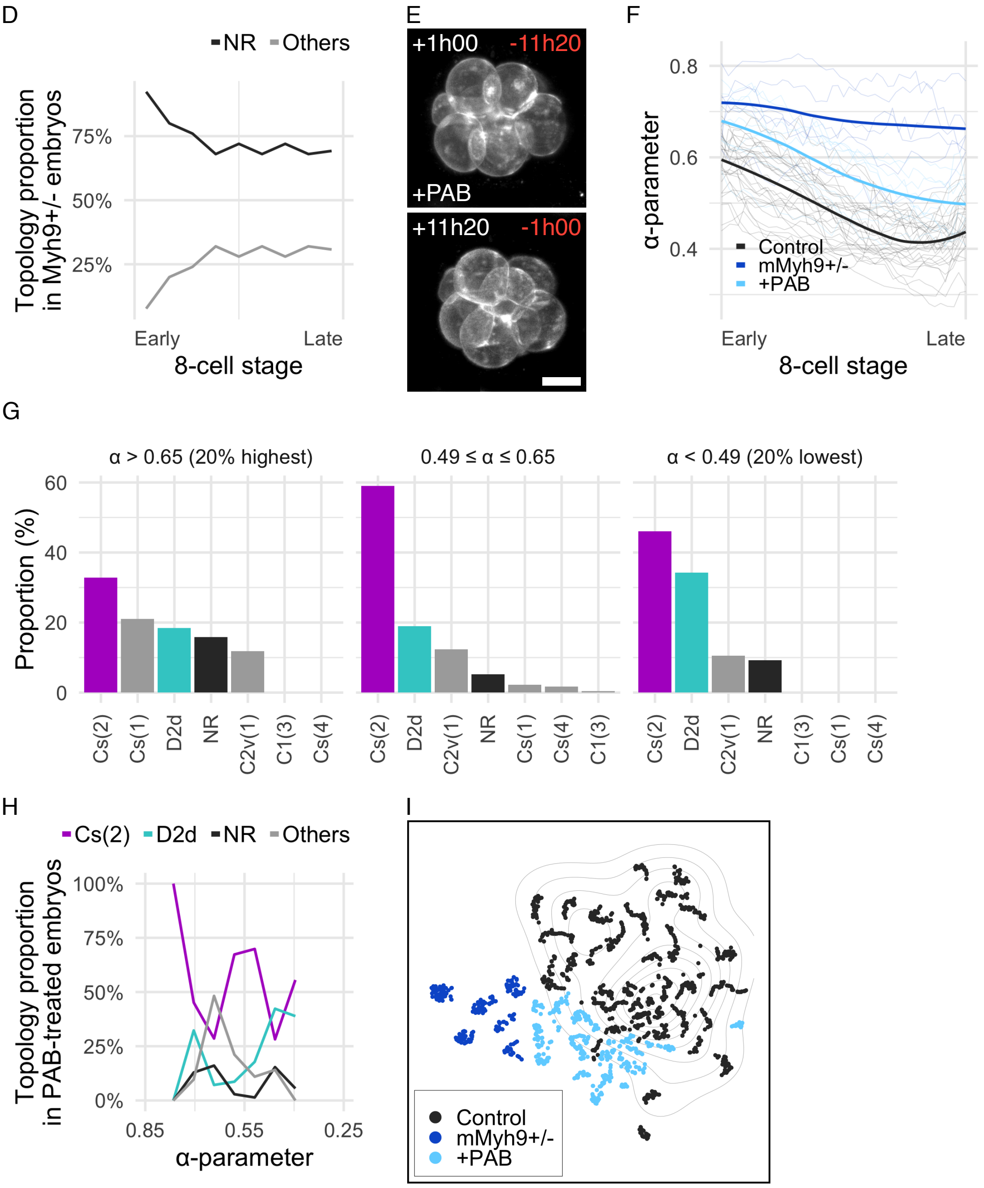

D
NR Others
Topology proportion in Myh9+/- embryos
75%
50%
25%
Early
Late
8-cell stage
E
+1h00 -11h20
+PAB
+11h20 -1h00
F
0.8
0.6
0.4
α-parameter
Control
mMyh9+/-
+PAB
Early
Late
8-cell stage
G
α > 0.65 (20% highest)
0.49 ≤ α ≤ 0.65
α < 0.49 (20% lowest)
Proportion (%)
60
40
20
0
Cs(2) Cs(1) D2d NR C2v(1) C1(3) Cs(4)
Cs(2) D2d C2v(1) NR Cs(1) Cs(4) C1(3)
Cs(2) D2d C2v(1) NR C1(3) Cs(1) Cs(4)
H
Cs(2) D2d NR Others
Topology proportion in PAB-treated embryos
100%
75%
50%
25%
0%
0.85
0.55
0.25
α-parameter
I
Control
mMyh9+/-
+PAB

**Figure S5. Compaction and surface contractility drive topological transitions.**
(**A**) Topological transition flow of mouse embryos, binned by the experimentally measured value of the $\alpha$-parameter (between $\alpha = 0.65$ and $\alpha = 0.35$), from and to $C_s(2)$ (magenta), $D_{2d}$ (cyan), non-rigid packings (NR, black) and other rigid packings (grey). Stream thickness represents the experimental proportion of topologies within each value of $\alpha$. (**B,C,G**) Distribution of the proportion of $C_s(2)$ (magenta), $D_{2d}$ (cyan), non-rigid packings (NR, black) or other minimally rigid packings (grey) identified in control (B, $n = 29$), mMhy9$^{+/-}$ (C, $n = 6$) and PAB treated (G, $n = 9$) mouse embryos for the 20% highest (left), the 60% middle values (middle) and the 20% lowest (right) $\alpha$-parameter.
(**D**) Evolution of the proportion of $C_s(2)$ (magenta, not present), $D_{2d}$ (cyan, not present), non-rigid (NR, black) or any other rigid packings (Others, grey) in mMyh9$^{+/-}$ mouse embryos as a function of the normalised progression through the 8-cell stage. $n = 6$ embryos. (**E**) Max projection of a representative live imaging of PAB treated embryos at the beginning (top) and the end (bottom) of the 8-cell stage ($n = 9$ embryos). Time in hours after the beginning (white, top-left corner) and before the end (red, top-right corner) of the 8-cell stage. Scale bar, 25 $\mu$m. See also Video S6. (**F**) Normalised time course through the 8-cell stage of the mean $\alpha$-parameter for control mouse embryos (black, $n = 29$), PAB treated embryos (light blue, $n = 9$) and mMyh9$^{+/-}$ mutants (blue, $n = 6$). Light colours, individual tracks. (**H**) Proportion of $C_s(2)$ (magenta), $D_{2d}$ (cyan), other rigid packings (Others, grey) and non-rigid packings (NR, black) as a function of the compaction parameter in PAB treated embryos ($n = 9$). (**I**) tSNE projection of the morphomap of control mouse embryos (black, $n = 29$), PAB treated embryos (light blue, $n = 9$) and mMyh9$^{+/-}$ mutants (blue, $n = 6$). Isolines, density map at the end of the 8-cell stage of control embryos.

### Figure S6

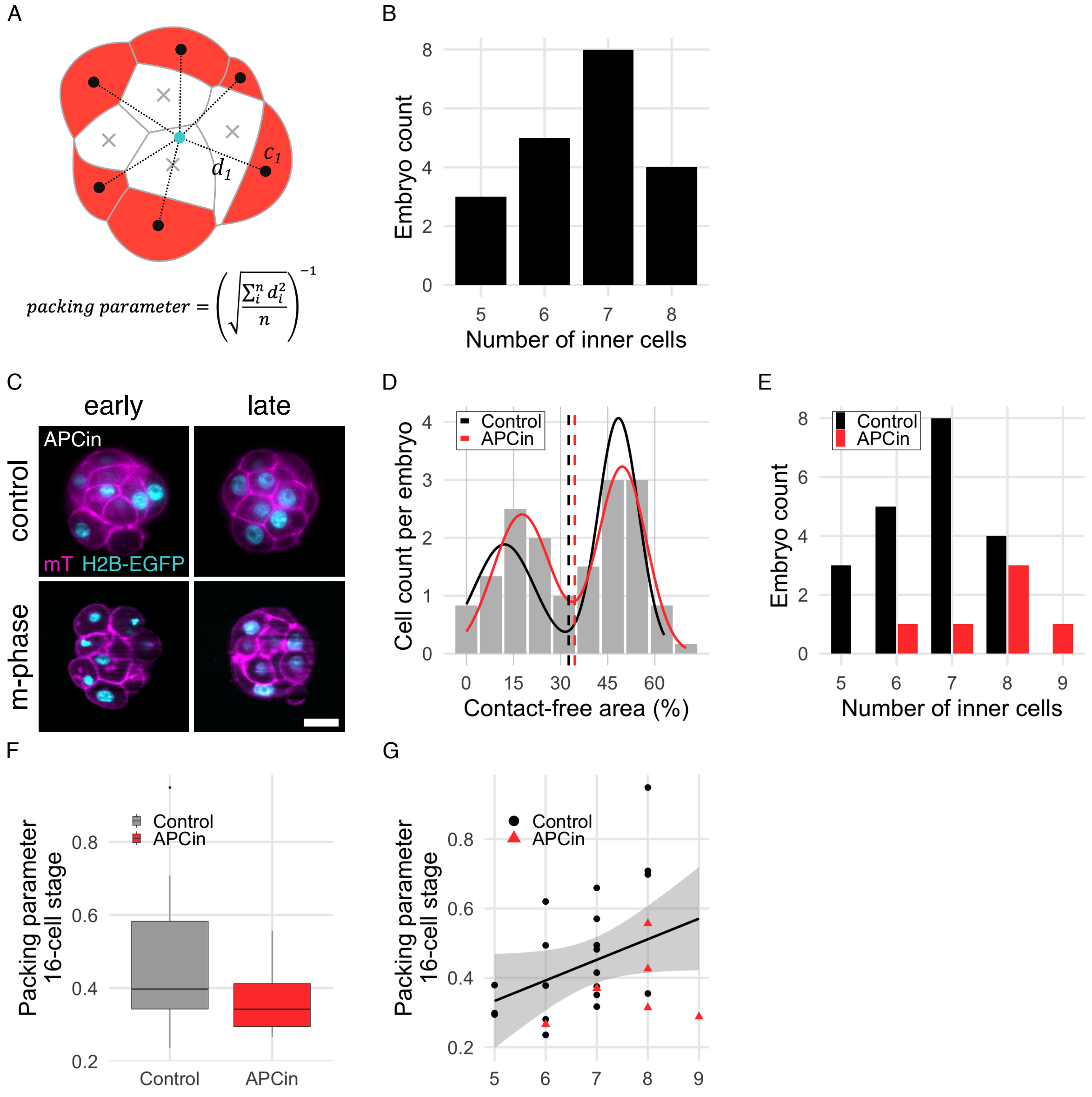

A
packing parameter = $\left(\sqrt{\frac{\sum_i^n d_i^2}{n}}\right)^{-1}$
$d_1$
$c_1$
B
Embryo count
Number of inner cells
C
early
late
control
m-phase
APCin
mT H2B-EGFP
D
Control
APCin
Cell count per embryo
Contact-free area (%)
E
Control
APCin
Embryo count
Number of inner cells
F
Control
APCin
Packing parameter
16-cell stage
G
Control
APCin
Packing parameter
16-cell stage

**Figure S6. Variability in cleavage timing promotes robustness in morphogenesis and ICM-TE patterning.**
(**A**) Visual representation of the packing parameter. Black dot, centre of mass of outer cells. Grey cross, centre of mass of inner cells. Cyan dot, centre of mass of the embryo. Dotted line, distance between outer cell centres and the centre of mass of the embryo, noted $d_i$. n, number of outer cells. (**B**) Distribution of the number of inner cells in 16-cell stage mouse embryos ($n = 20$). (**C**) Max projection of a representative live imaging of embryos from group control (first row) and group m-phase (second row, treated with APCin, $n = 6$ embryos) one hour after the beginning (left column) or before the end (right column) of the 16-cell stage. Magenta, mT (membranes). Cyan, H2B-EGFP. Scale bar, 25 $\mu$m. (**D**) Bi-modal count distribution of the contact-free area of individual cells from embryos in group m-phase treated with APCin (grey bars, $n = 96$ cells pooled from 6 embryos) fitted with the sum of an inner and an outer gaussian distribution (red line, $\mathrm{mean} \pm \mathrm{s.d.} = 17.0\% \pm 8.9$ and $50.0\% \pm 7.6$ respectively). Black line, distribution of the contact-free area of individual cells in group control, for comparison. Red dashed line, cut-off between inner and outer cells, defined as the intersection point of the two gaussians (cut-off = 34.5%). Black dashed line, cut-off in group control. (**E**) Distribution of the number of inner cells in 16-cell stage mouse embryos from the group control (black, $n = 20$) and the group m-phase treated with APCin (red, $n = 6$). (**F**) Packing parameter at the 16-cell stage in group control (grey, $n = 20$) and group m-phase treated with APCin (red, $n = 6$). (**G**) Packing parameter as a function of the number of inner cells in group control (black, $n = 20$) and group m-phase treated with APCin (red, $n = 6$). Pearson correlation $R = 0.369$ ($P = 0.06$). Solid line, linear regression. Shaded ribbon, standard error.

#### Supplementary Videos

**Video S1. Live imaging and segmentation of a mouse embryo.**
(**Left**) Maximum projection of live the imaging of a mouse embryo expressing mT (magenta); H2B-EGFP (cyan) developing from the 4- to the 16-cell stage. Time in hours post imaging (hh:mm:ss). (**Right**) Cell segmentation of the same embryo at the 8-cell stage. Segmentation before and after the 8-cell stage inter-mitotic period is not shown. Each cell is represented with a different colour.

**Video S2. Live imaging and segmentation of a monkey embryo.**
(**Left**) Maximum projection of the live imaging of a monkey embryo micro-injected with a single mRNA encoding memb-tdTomato (magenta) and H2B-EGFP (cyan) fused with self-cleaving 2A peptide, developing from the 4- to the 16-cell stage. Time in hours post imaging (hh:mm:ss). (**Right**) Cell segmentation of the same embryo at the 8-cell stage. Segmentation before and after the 8-cell stage inter-mitotic period is not shown. Each cell is represented with a different colour.

**Video S3. Live imaging and segmentation of a rabbit embryo.**
(**Left**) Maximum projection of the live imaging of a rabbit embryo micro-injected with two mRNAs encoding myrTagRFP-T (magenta) and H2B-EGFP (cyan), developing from the 4- to the 16-cell stage. Time in hours post imaging (hh:mm:ss). (**Right**) Cell segmentation of the same embryo at the 8-cell stage. Segmentation before and after the 8-cell stage inter-mitotic period is not shown. Each cell is represented with a different colour.

**Video S4. Timelapse of a topological transition from $C_1$(3) to $D_{2d}$.**
The red and the blue cells are initially not in contact ($C_1$(3), 5:30), then transiently ($C_s$5, 7:30), then established ($D_{2d}$, 8:45) and strengthen ($D_{2d}$, 11:30). Time post imaging (hh:mm:ss).

**Video S5. Max projection of the live imaging of a representative mMyh9+/- embryo developing from the 4- to the 32-cell stage.**

**Video S6. Max projection of the live imaging of a representative PAB treated embryo developing during the 8-cell stage.**

#### Supplementary Tables

**Table S1. Imaging parameters of the live-imaged embryos used in this study.**
The leading letter in embryos' name corresponds to the experiment, therefor indicates embryos imaged at the same time. ∂xy, lateral resolution. ∂z, step size between consecutive optical slices as estimated after correction due to a technical problem affecting experiments A, B, C, D, E and H (see Methods). For consistency, correction was applied to every embryo. Embryos with uncorrected step size were not used in spatial analyses. ∂t, time difference between the beginning of two consecutive stacks of optical slices. Duration corresponds to the period of acquisition.

**Table S2. Use of live imaging dataset in figures.**
The leading letter in embryos' name corresponds to the experiment, therefor indicates embryos imaged at the same time.
